# FaPDA: Facial paralysis detection algorithm applied in mice

**DOI:** 10.1101/2024.09.02.610902

**Authors:** Elías Perrusquia Hernández, Diego Israel Villeda Arias, Claudia Daniela Montes Ángeles, Rey David Andrade González, Joel Lomelí González, Isaac Obed Pérez-Martínez

## Abstract

Facial paralysis is characterized by an injury to the facial nerve, causing the loss of the functions of the structures that it innervates and changes in the central nervous system in the motor cortex. Therefore, it is important to use experimental models to further the neurophysiological study. Currently, the models have some limitations for the study of facial paralysis. Therefore, the development of an algorithm capable of automatically detecting facial paralysis and overcoming the existing limitations is proposed. C57/BL6 mice were used, which produced irreversible facial paralysis with permanent effects and reversible facial paralysis, where the effects of the paralysis disappear within the first 15 days after the nerve injury. Video recordings were made of the faces of paralyzed mice to develop the algorithm for detecting facial paralysis applied to mice, which allows us to detect the presence of reversible and irreversible facial paralysis automatically. At the same time, the algorithm was used to track facial movement during oral stimulation with sucrose and extracellular electrophysiological recordings in the anterolateral motor cortex. In the basal state, mice can make facial expressions associated with a pleasurable state, and the algorithm detects this movement; at the same time, the movement correlates with the activation in the cortical area. In the presence of facial paralysis, the algorithm can not detect movement. Therefore, it concludes that the condition exists, and the neuronal activity in the cortex is affected with respect to the evolution of facial paralysis. Therefore, the facial paralysis detection algorithm applied to mice allows deduce the presence of experimental facial paralysis, in the presence of oral stimulation or without it, at the same time that cortical electrophysiological recordings can be made for the neurophysiological study of facial paralysis.

**Significance Statement:** Automatic detection of facial paralysis by video recording of the face of mice can help to identify the presence of the condition more easily. It can also used to study facial movement, like facial expressions, and their neural correlates in cortical and subcortical strata. This will allow understanding in depth the level of affectation that facial paralysis can produce at the peripheral and central nervous system levels.

## Introduction

Facial paralysis is a condition where the facial nerve is damaged and results in the loss of function of the structures it innervates (Maspero, Farronato, Guenza, & Farronato, 2017; Walker, Mistry, & Mazzoni, 2024). It has been reported that 30% of those who have facial paralysis experience long-term effects, and 5% present a high degree of sequelae after apparent recovery (Maspero et al., 2017). Damages are not only limited to the muscles or other facial structures. They have also been observed in the central nervous system, in structures related to the motor and sensory response, such as the primary motor and somatosensory cortex (Klingner et al., 2011; Song et al., 2017).

Therefore, different experimental strategies have been proposed for the study of facial paralysis, mainly in rodents (rats and mice). The cardinal signs that have been used to recognize the presence of experimental facial paralysis are the movement of the vibrissae of the face, the eye area, and the nose orientation (Severson, Xu, Yang, & O’Connor, 2019; Sugita et al., 1995; Zeng et al., 2024). However, in order to apply these strategies, the use of specialized systems is required, as well as specific preparation of study subjects and movement restriction (Attiah, de Vries, Richardson, & Lucas, 2017; Severson et al., 2019; Vajtay et al., 2019).

The conditions mentioned earlier represent a limitation for the study of facial paralysis and its effects on the central nervous system, which are currently shown as electrophysiological changes in action potentials and dendritic retraction in the motor cortex and corticofacial structures (Munera, Cuestas, & Troncoso, 2012; Urrego, Munera, & Troncoso, 2011). Nevertheless, a facial movement begins in premotor structures, which could be affected by facial paralysis, such as the anterolateral motor cortex (ALM), which is responsible for the planification, coordination, and execution of movements (N. Li, Chen, Guo, Gerfen, & Svoboda, 2015). It has been related to the movement of facial muscles (Haiss & Schwarz, 2005).

Hence, the present work attempts to use a new strategy that allows the study of facial paralysis and its conditions in the central nervous system. The use of image processing is proposed for the automatic detection of facial paralysis since, in recent years, it has been used to track facial movement by detecting vibrissae on the mouse face (Severson et al., 2019) to give electrophysiological tracking of orofacial movement (Syeda et al., 2024) and to detect, quantify and classify emotional facial expressions (Dolensek, Gehrlach, Klein, & Gogolla, 2020; Le Moene & Larsson, 2023; Tanaka, Nakata, Hibino, Nishiyama, & Ino, 2023). These tools have resulted in great support for the study of orofacial movements and their participation at the level of the central nervous system (CNS).

In this sense, two models of facial paralysis (transection or compression of facial nerve) were used for the design, creation, and validation of an algorithm for the detection of facial paralysis applied in mice (FaPDA). This allowed us to determine if the study subject was healthy, paralyzed or with an apparent recovery from the condition. FaPDA was designed to be used in conditions of semi-restricted movement, with and without stimuli, during the evaluation and monitoring of facial expressions, performing extracellular electrophysiological recordings at the same time.

Our results show that facial transection injury generates irreversible facial paralysis. In contrast, compression injury generates reversible paralysis, showing disorders in whisker and facial movements in the middle and anterior part of the mouse’s face. These two areas were used for the development of the FaPDA, which allowed us to determine the presence of both reversible and irreversible facial paralysis. Similarly, it was found to be useful for tracking facial movement during the study of facial expressions and their neural correlates in ALM. Finally, facial paralysis causes an inability to make pleasurable facial expressions and a decrease in firing frequency in ALM neurons in the irreversible model. On the other hand, in the reversible model, there is an apparent recovery of the ability to make facial expressions. Still, the neuronal activity in ALM is not comparable with baseline values.

Therefore, PaPDA is a viable proposal for the study of experimental facial paralysis, with or without the presence of sensory stimuli, for orofacial movement studies and during electrophysiological recordings at cortex level.

## Methodology

### Animals

Twelve male and female C57BL/6 mice were used, aged 2 to 6 months. All procedures were carried out in accordance with the standards of the ethics committee of the Iztacala School of Higher Studies of the National Autonomous University of Mexico (CE/FESI/012022/1469). All subjects were housed in polycarbonate boxes. The housing room had a temperature of 25° C, under a 12-12 hour light-dark cycle (starting at 8:00 am). Subjects were provided with standard food and water ad libitum. Only during conditioning and testing were they deprived of water, allowing a maximum consumption of 4 ml per day.

### Surgery for head-fixation device implantation and craniotomy

Mice were anesthetized with isoflurane (3% for induction, 1% for maintenance). Under the surgical plane, an incision was made on the scalp, and tissues were removed to expose the skull. Using a stereotaxic system, ALM was located 2.5 mm anterior and 1.5 mm lateral to bregma. Two mice underwent a rectangular craniotomy with a 2 × 2 mm extension centered on the location of ALM. The craniotomy opening was covered with dental silicone, and tissues were repositioned. All subjects had a device (made of polylactic acid) for the head-fixation system placed on the back of the skull, fixed with cyanoacrylate. Mice were given seven days for recovery.

### Facial nerve surgery

Mice were anesthetized following the protocol mentioned above. Under the surgical plane, an incision was made posterior to the right ear of the mice. Tissues were dissected until the trunk of the facial nerve was observed. Finally, one group of mice had their nerve compressed with dissection forceps twice for 30 seconds with an interval of 10 seconds between compressions; the force was recorded with an Arduino-compatible compression sensor (Walfront, Model: Walfront9snmyvxw25) (compression injury model). The next group underwent a complete cut of the facial nerve (transection injury model). Tissues were repositioned, and the incision was sutured. For sham group, the facial nerve was only observed, and the incision was sutured.

### System

For whisker movement and facial expressions recording, as well as acute neural recordings, a semi-restricted movement system was used. The system consisted of a 32 cm diameter ball held by two lateral axes to an acrylic base, which allowed it only to move forward or backward. On the ball, there was a fixing bar where the attachments previously implanted in the head of the mice fit.

Video recording was made with a 30Hz acquisition and 2K resolution cell phone camera (Blackview BV8800). The camera was placed perpendicular to the right face of the mouse. Video recording of the whisker was made with a 120Hz acquisition and HD resolution cell phone camera (Xiaomi Note 8). The camera was placed on the superior part of the mouse’s face and allowed the recording of the right and left whiskers.

### Facial paralysis model

Nine mice were used: 3 with facial nerve transection, 3 with compression, and 3 sham mice. Twenty-three measurements were made starting with day -1 (baseline), at .5, 6, 24 hours after the injury, and finally, from day 2 to 20 after the injury. Whisker movement was measured, without stimulation, through the angulations formed by the vibrissae with its insertion in the whiskerpad, the initial position of the whisker was considered as 0°. From the initial position, subsequent angulations were obtained, forming a continuous signal. The minimum angulations were located throughout the signal. From each minimum to the minimum value, a whisker movement cycle was considered. This consisted of protraction movement, moving from the most posterior position of the whisker (minimum angle) to the most anterior (maximum angle), and retraction, moving in the opposite direction to the previous movement. Each movement cycle respected a certain amplitude, which was obtained by subtracting the minimum angle from the maximum. The average amplitude of all the cycles in 2 seconds was taken as the threshold. Above this threshold, it was considered as a long amplitude, and below this threshold, it was considered as a short amplitude. Each movement cycle was considered one Hz, and all the movement cycles throughout the 2 minutes of the video recording were used.

### Whisker movement tracking

For the study of whisker movement, acrylic paint was used to mark a whisker from both the ipsi- and contra-lateral sides of the lesion. Video recording was transformed into frames and processed with MATLAB®. The initial step of processing was to identify the color of the whisker painted in each frame. Then, the identity of the pixels with this color was obtained. The image was binarized, where the pixels belonging to the whisker were replaced by white, whereas the rest were black. MATLAB Steel function was used to clean the detected area, which allowed us to eliminate pixels outside the required area (within a range of 5 pixels around). Finally, with the angle function, the angulation of the whisker was obtained, taking into account the reference point indicated and previously described. The smallest angle detected during the video was always considered 0°.

### Facial motion analysis

To study facial expressions and movement, the histogram-oriented gradient (HOG) image descriptor was used in each frame using the following parameters: oriented histogram content = 8, cell box length size = 32 pixels, and cell per block = 1, along with a previous normalization processing using the power compression law. For the study of facial paralysis, mice face image was cut into three zones: anterior, middle, and posterior zone. This was done over 2 minutes of evaluation. Three thousand six hundred frames were obtained per zone for the study of facial expressions. An intrinsic state (pleasure and disgust) and a neutral state (water) were studied five seconds before (control) and after oral stimulation were analyzed, which corresponded to a total of 300 frames. Each frame was transformed from RGB to grayscale, and the mathematical vectors (HOGs) were subsequently obtained. All videos were processed with MATLAB®.

### Facial paralysis detection algorithm applied in mice (FaPDA) design

The computational algorithm for detecting facial paralysis is a bimodal decision system, which consists of indicating whether the condition exists or not. To do this, the movement threshold of different sections of the face (anterior, middle, and posterior areas) is taken into account. The threshold is obtained from mice with facial paralysis by transection and is determined by the sum of the average of the difference between HOGs and their standard deviation.

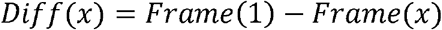

Diff= Difference between frames

Frame= Frame converted into HOG mathematical vector.

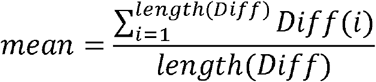

mean= average of differences between frames.

i= evaluations (23, including baseline)

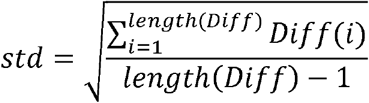

std= standard deviation of the difference between frames.

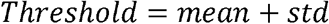

To determine the presence of facial paralysis, different variables were taken into account: areas of the face, average threshold per group or individual, and the normalization of the data. In this way, 9 facial paralysis decision models were obtained. To decide if there is facial paralysis, there must be values lower than the threshold of the different areas. If this occurs, the value is determined with a binary value of 1. Otherwise, the value is 0.

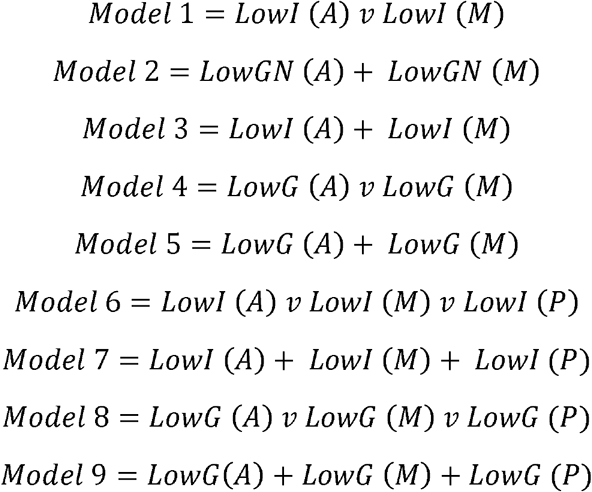

LowG= Binary values 1 (below threshold), 0 (above threshold), where the threshold is averaged per group.

LowGN= Binary values 1 (below threshold), 0 (above threshold), where the threshold is averaged per group, data to obtain threshold and binary values are normalized.

LowI= Binary values 1 (below threshold), 0 (above threshold), where threshold is individual per mouse.

A= anterior zone

M= middle zone

P= posterior zone

N= mice per group (n=3)

Subsequently, an attempt was made to extract the threshold individually for mice injured by transection and compression. To do this, mice injured by transection were used, where the threshold for each mouse was obtained, taking into account the average of the difference between frames.

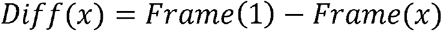

Diff= Difference between frames.

Frame= Frame converted into HOG mathematical vector.

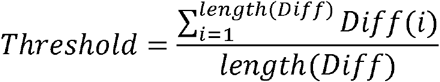

Threshold= Average of the differences between frames. i= evaluations (23, including the baseline)

Once the threshold was created, the number of evaluations for the optimal functioning of the algorithm was sought.

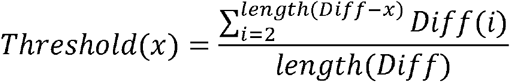

x= number of tests (up to 22).

It was determined that two assessments, one at baseline and one at a standstill, were sufficient to obtain an appropriate threshold. Finally, the assessment that would be ideal for the functioning of the algorithm was sought.

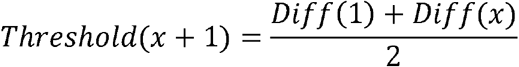

This technique is used for mice individually for the transection and compression injury group with the previously used model.

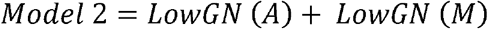

As a last resort, the algorithm was used to track movement in real-time.

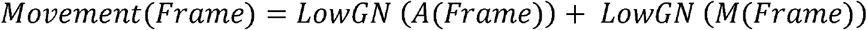

Movement = movement detected by the algorithm. If the value is less than 2, there is movement; otherwise, there is no movement.

If the percentage of moments without movement is greater than 90%, the system concludes that the mouse is paralyzed.

### Oral stimulation test

Behavior assay consisted of the release of sucrose (20%), quinine (0.3 mM), and regular water directly into the mouth of the mice through an intraoral cannula. Mice were placed in the semi-restricted head system, recording the walk (speed at which the mouse moved on the system) and facial expressions. A 5 minute-baseline recording was made, in which no stimulus was presented. Further, the video recording was made during oral stimulation. For the release of a solution, mice must avoid walking for 10 seconds. When this condition was met, a solution semi-randomly chosen was delivered, 8 drops of 4 microliters each, and facial video recording was made for 1 minute. This was considered a trial. For a new trial to start, the condition of not walking for 10 seconds had to be met, and a session consisted of 5 trials per solution. Finally, a 5-minute video recording was made with the solution infusion pump activated without releasing any solution. For the electrophysiological recordings, the previous protocol was repeated. However, stimulation was performed only with sucrose.

### Electrophysiology

A 16-channel microelectrode array (Tucker Davis Technologies, OMN1030-16) was acutely implanted in CMAL (Figure 1). Microelectrode voltage signals were acquired, digitalized, and filtered with an Open Ephys multichannel acquisition processor. Only individual neurons with action potentials with signal-to-noise ratios greater than 3:1 were selected for analysis and sorted using the Plexon Offline Spike-Sorter system.

**Figure 1.**
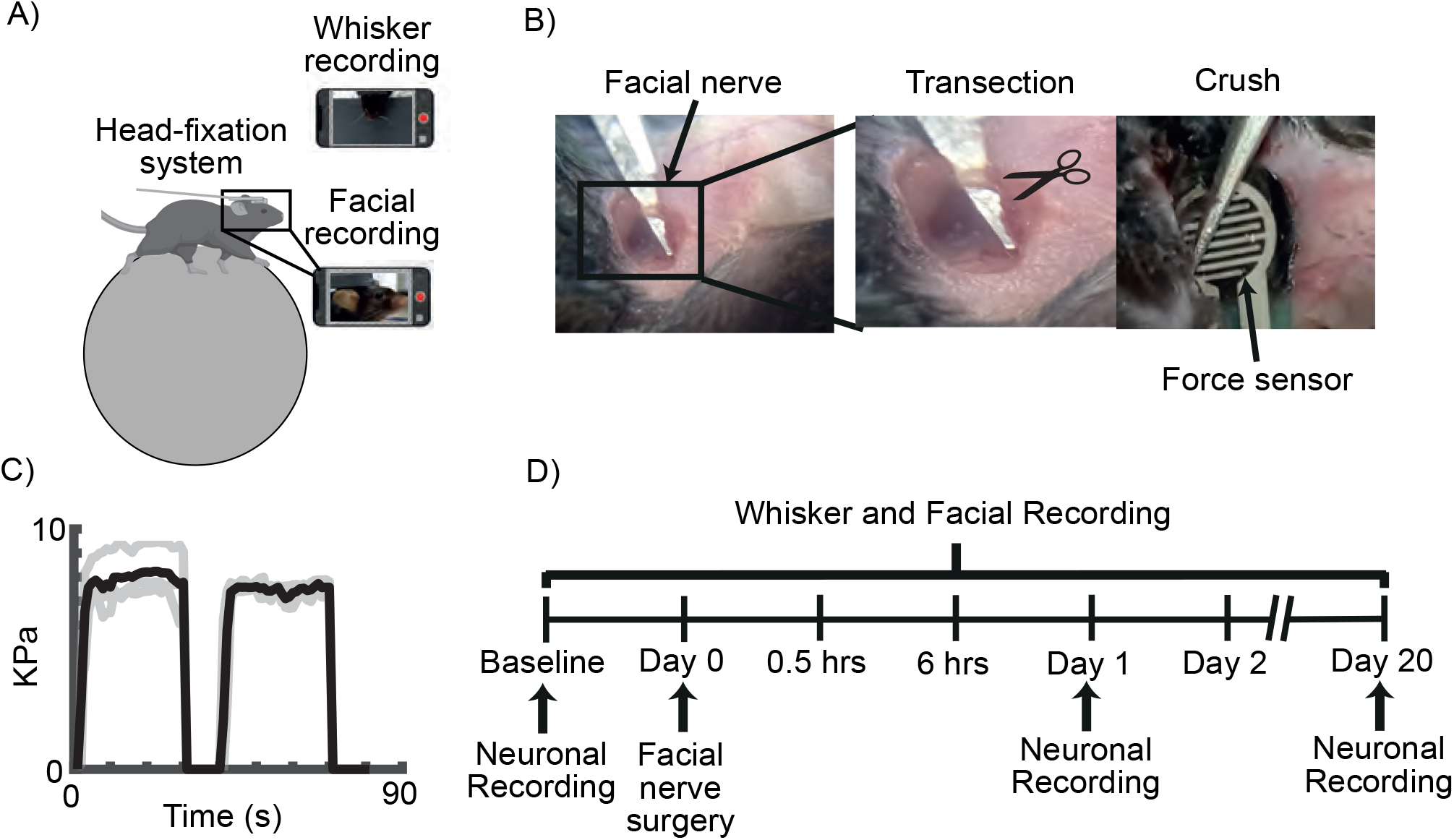
Experimental design. A) Schematic of the semi-restricted movement system and simultaneous video recording of whiskers and face of the mouse. B) Surgical process of facial paralysis models of nerve injury by transection and compression. C) Monitoring of the force applied to the facial nerve when it is compressed, the grey lines represent each mouse (n=3) and the black line its average (p>0.05 ANOVA-one way test, Figure 1-1). D) Experimental timeline. Detailed statistics in Extended Data Table 1-1

### Creation of the facial prototype

First, a basal or resting state facial prototype was determined, which was created using frames during the period prior to oral stimulation (250 frames), transformed into HOGs, and averaged. This basal prototype in HOGs was compared with the entire stimulation set with 20% sucrose, quinine, and water, calculating the Pearson correlation coefficient. From each solution, the 10 frames with the greatest difference with the basal facial prototype (those with the lowest correlation coefficient) were taken. The average of these frames resulted in a single HOG considered as the emotional prototype (pleasant for sucrose, disgust for quinine, and neutral for water).

### Statistical analysis

The comparison between the values obtained by the force sensor when compressing the facial nerve was made with a one-way ANOVA analysis, comparing the applied force between the 3 subjects in the group. During the tracking of the whisker movement, ischange function was used to detect the moment with the greatest change in a signal, which includes the following values.

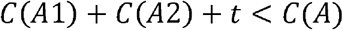

A=data vector (whisker movement signal)

A1=first segment of the vector

A2=second segment of the vector

C=cost function

t=threshold created by the MATLAB threshold function

The threshold function creates an object of discrete state threshold transitions and is specified by the mean transition levels. The cost function tells us how close a segment is to this segment.

After that, the amplitudes before and after the change point were taken into account and were analyzed with Student t-test analysis. The area under the curve of the whisker movement cycle of the 23 evaluations performed (baseline, 30 minutes, 6 hours, and from day 1 to 20 after the facial injury) was obtained and analyzed with one-way ANOVA and a Tukey post hoc test. To study the proportion of long and short whisker movement amplitudes, a chi-square goodness-of-fit analysis was used. Whisker movement frequency values were organized and graphed using the MATLAB spectrogram function.

During the tracking of mouse facial movement in the three facial areas, a one-way ANOVA analysis with a Tukey post hoc test was used. A Pearson correlation analysis was applied between the whisker movement amplitudes and the difference between the HOG of the mice’s faces. For the analysis of facial expression, the frames before and after the release of the different solutions were used; using the MATLAB cluster-gram function, an automatic hierarchical tree clustering was performed. A t-distribution stochastic neighbor analysis (t-SNE) was performed for the clustering of frames associated with a particular solution. Finally, a Pearson correlation analysis was performed between the facial prototype (previously described) and the frames before and during oral stimulation.

To obtain the accuracy of the system, the value d’ was used, which considers:

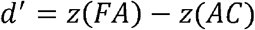

AC=Hit (the algorithm detects facial paralysis after transection facial injury)

FA=False alarm (there is no facial paralysis, and the algorithm does detect it)

z=typical z value

For the electrophysiological analysis, a Wilcoxon rank sum analysis was performed for the population neuronal activity with data before and after oral stimulation with sucrose, activation of the solution infusion pump, and at the time of starting to walk. The neuronal activity consists of the average of the z value (normalized in a range of -2 to 2) obtained from the peristimulus time histogram (PSTH), which shows us the amount of activation of the neurons recorded in an interval of (100 ms). The z value was obtained with the following formula:

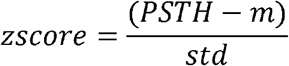

PSTH=neuronal activity

m=mean obtained from the neuronal activity prior to stimulation (10 s).

std=standard deviation obtained from the neuronal activity prior to stimulation (10 s).

Finally, a cross-variance analysis was used for the relationship between neuronal activity, facial expression and facial movement. The cross-variance values between neuronal activity with facial expression and neuronal activity with facial movement were analyzed with Pearson correlation. For more details on the p values of each statistical test, see attached Tables.

### Code accessibility

The code/software described in the article is open source and is freely available on GitHub (https://github.com/Elias4444/FaPDA.git). The code is also available in Extended Data 1. FaPDA can be used in MATLAB software (version 2023b with image processing toolbox). The specification of a computer is anyone that can run MATLAB software.

## Results

### Reversible and irreversible facial paralysis model in mice

To determine the presence of facial paralysis in the study subjects, a semi-restricted movement system and bilateral video recording of their whiskers were used (Figure 1A). Two models of facial paralysis were used: transection and compression. In addition to these, a sham group was added as a control group (Figure 1B). In order for the transection injury to be uniformly performed in all subjects, a complete cut of the nervous tissue was made, and the loss of continuity of the facial nerve was confirmed. In the compression injury, a force sensor was used (see materials and methods) which measured the applied pressure on the nervous tissue (Figure 1B). This way, the measurements obtained in each subject did not present statistically significant differences (Figure 1C), which indicated that the injuries were comparable with each other.

To study the effects of facial paralysis, video recording of whisker movement was used. A whisker was tracked both ipsi and contralateral to the lesion. Its position was determined in each frame of the video. The position was given by the angle formed between the whisker and its insertion point in the whisker pad. The accumulation of the whisker positions throughout the video recording represented a signal. Using a mathematical model (Matlab function: ischange), which allowed us to determine abrupt changes in a signal, we searched for the frame where the whisker had the greatest changes in its angulations, which we called the change point. (Figure 2-1A and B). Minimum and maximum peaks of the signal were detected two seconds before and after the change point (Figure 2-1B); this was used to determine the amplitude of the whisker movement, which consisted of the subtraction between the angle of the most anterior position (minimum peak) and the posterior position (maximum peak). With the above, a difference can be observed between the amplitudes of the movement before and after the point of change, prior to the facial injury for the transection, compression, and sham group (Figure 2-1C). This showed us the change between resting state of the whisker and the moment in which movement began in all groups. In this way, only the moments in which there was whisker movement were analyzed.

Whisker movement cycle, given by the protraction movement (travel from the most posterior position to the most anterior position of the mouse’s face) and retraction (travel from the most anterior position to the most posterior position of the mouse’s face), was used to study the degree of mobility that whiskers had. The area under the curve of this cycle was used to determine the movement width, both before and after nerve injury (Figure 2). During baseline evaluation, the 3 groups (transection, compression, and sham) behaved in a similar way, where the whiskers respected retraction and protraction movement (Figure 2A-C). Following nerve injury by transection, whisker movement cycle was altered, showing a decrease in the amplitude of movement from half an hour to day 20 after the injury (Figure 2A and 3A). The compression injury showed similar effects to those of transection during the first 9 days. Subsequently, an increase in the amplitudes of the whisker movement cycle was observed until reaching baseline values from day 15 to day 20 after the injury (Figure 2B and 3D). The sham group maintained the amplitudes of its movement cycles throughout the evaluations, showing a decrease in their amplitudes on days 8 and 17 after surgery (Figure 2C and 3G). The opposite occurred on the contralateral side of the injury: in the transection and compression group, the whisker movement cycle or its amplitudes were not lost (Figure 2-2). This showed that the transection injury generates irreversible facial paralysis since the whisker mobility was never recovered. Compression injury is reversible since an apparent recovery of function is observed from day 15, and this is due directly to damage to the facial nerve and not to tissue manipulation.

**Figure 2.**
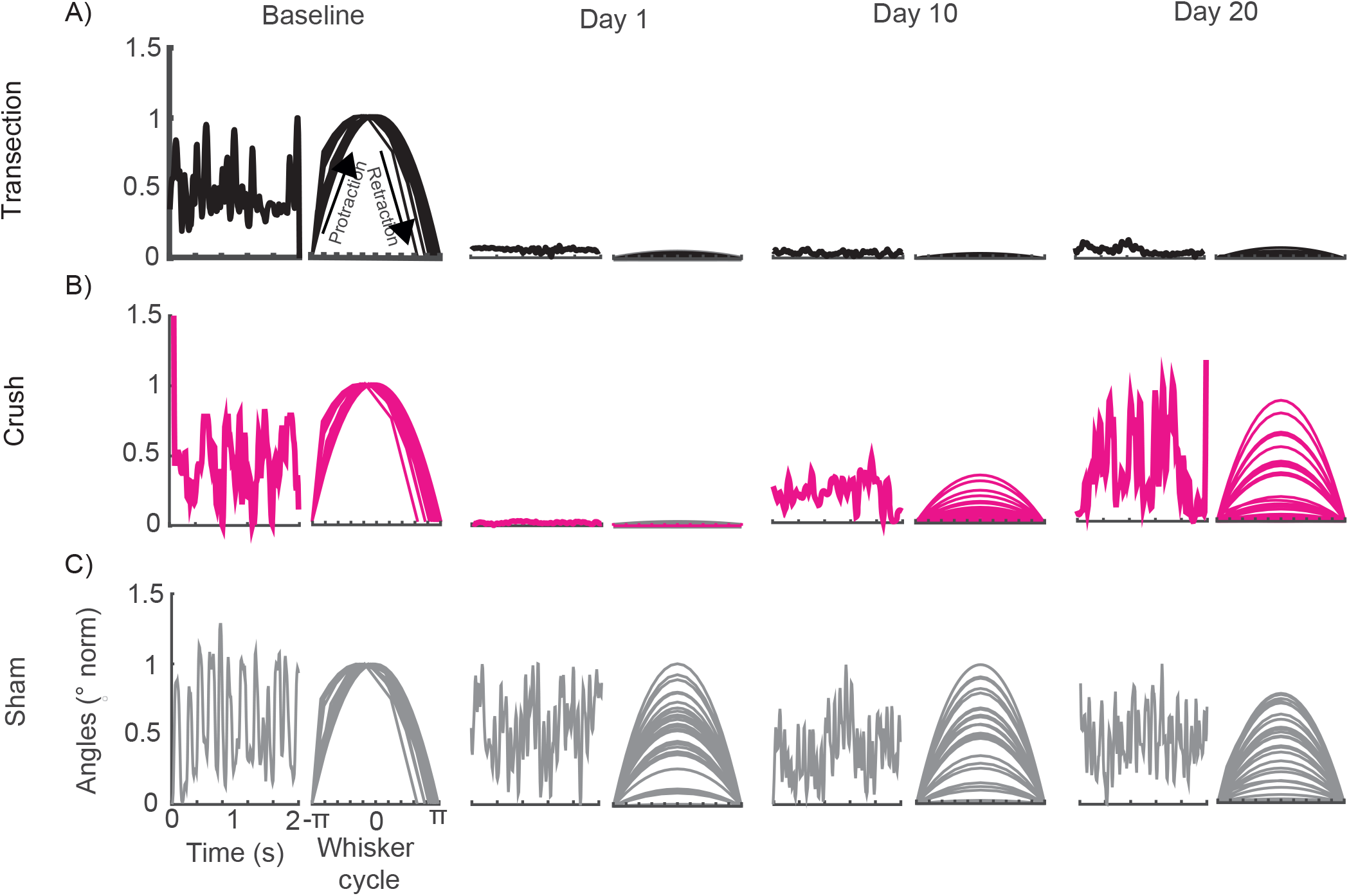
Effect of facial paralysis on whisker movement cycle. Whisker movement monitoring of a single mouse per group (A) transection, B) crush, and C) sham, prior to surgery and on days 1, 10, and 20 postoperatively. The graphs on the left of each column represent the dynamics of whisker movement over 2 seconds of evaluation, and on the right, their movement cycle is shown.

**Figure 3.**
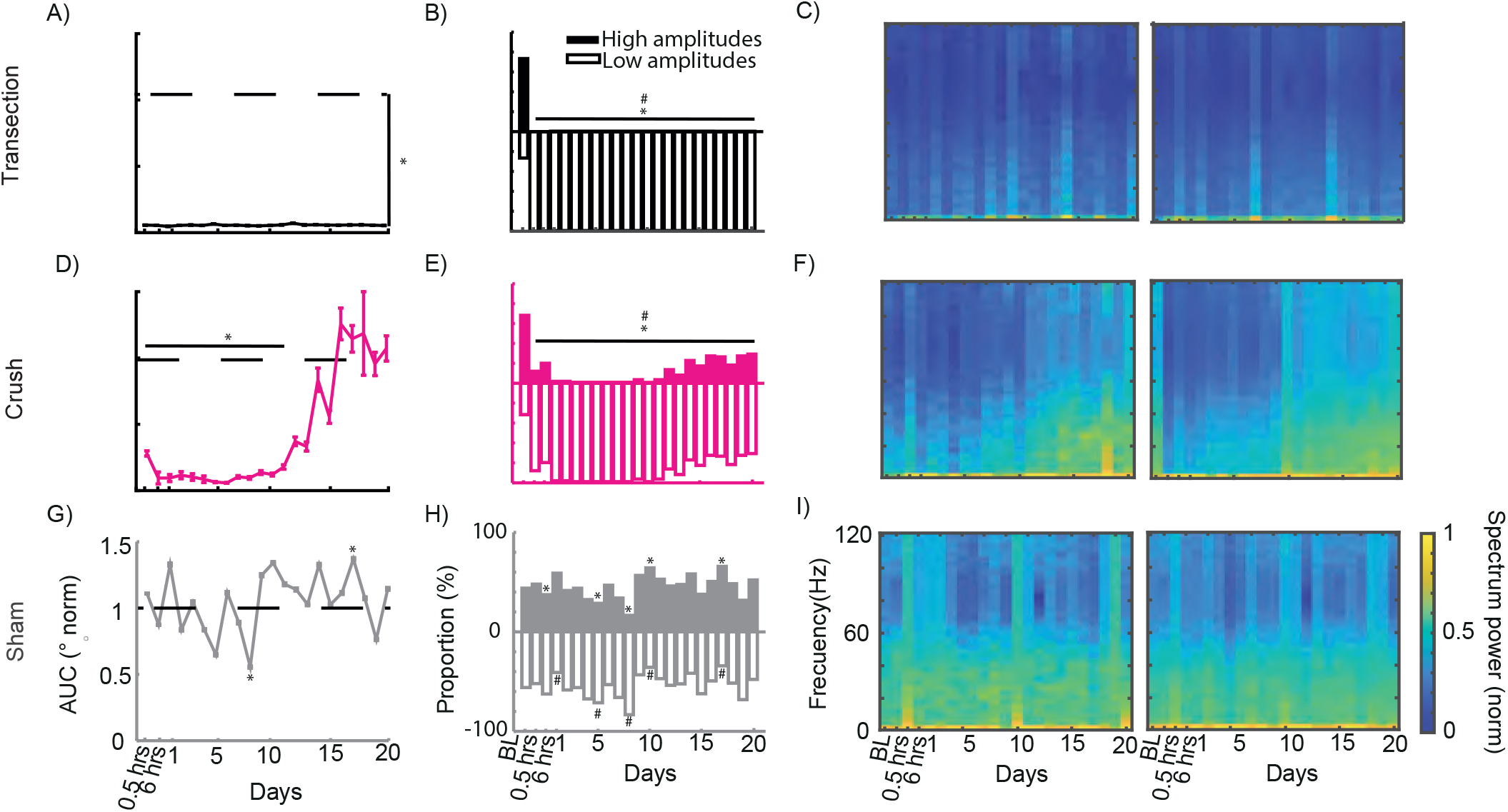
Physiological changes in whisker movement after facial paralysis. A) Area under the curve of the whisker movement cycle assessed from half an hour to day 20 after surgery (Baseline vs Lesion * p<0. 05 ANOVA one way test, Tukey test p<0.05, n=3). B) Proportion of high and low amplitudes obtained from the whisker movement cycle (Baseline vs Lesion in High Amplitudes * p<0. 05 and Baseline vs Lesion in Low Amplitudes # p<0. 05 Chi-squared test). C) Spectrogram of whisker movement frequency over 2 seconds, the left graph represents one mouse and the right the average of the transection injury group. This description is repeated in D (Baseline vs. Lesion * p < 0.05 ANOVA one-way test, Tukey test p < 0.05), E (Baseline vs. Lesion in High Amplitudes ** p < 0. 05 and Baseline vs. Lesion in Low Amplitudes ## p < 0. 05 Chi-squared test) and F for the compression injury group (n = 3); and G (Baseline vs. Surgery * p < 0.05 ANOVA one way test, Tukey test p < 0.05), H (Baseline vs Lesion in High Amplitudes * p < 0.05 and Baseline vs Lesion in Low Amplitudes # p < 0. 05 Chi-squared test) and I for the sham group (n = 3). The dotted lines in A, D, and G represent the baseline. Detailed statistics in Extended Data Table 3-1, Table 3-2, Table 3-3 and Table 3-4

### Physiological differences in whisker movement between transection and compression facial paralysis models

The first physiological aspect to be studied was the movement amplitude, which showed us the range of movement that the whisker is capable of. In transection facial injury, there was no wide movement of the whisker, and this was observed throughout the 20 days of assessment (Figure 3A). However, whisker movement did not have a single amplitude; therefore, all amplitudes were averaged over 2 seconds of evaluation. Any amplitude above the average was considered a long amplitude and below a short amplitude. Prior to the injury, long and short amplitudes are shown, which changed their distribution after the injury, where short amplitudes predominated (Figure 3B). Finally, the frequency of movement was obtained, where the prevalence of low frequencies (0-5 Hz) after the facial injury was observed (Figure 3C). This showed that irreversible facial paralysis not only causes a loss in the dynamics and range of whisker movement but also affects the frequency at which it is performed.

Compression injury showed a decrease in the range of movement during the first 11 days after the injury. From day 12, a recovery that reached basal values was seen (Figure 3D). As in the irreversible injury, prior to the nerve injury, long and short amplitudes are shown; subsequently, an increase in low amplitudes was observed until day 11. For day 12 and up to day 20, a progressive increase in high amplitudes was observed. However, at no time do they get comparable with basal state (Figure 3E). When analyzing the frequencies, it was observed that low frequencies predominate during the first 10 days after the injury (0-5 Hz). From day 11, a variation between 0 and 45 Hz is observed (Figure 3F). Thus, despite an apparent recovery in frequency and range of motion, there was difficulty in making wide movements of the whisker.

Finally, the sham group never loses the range of motion after surgery (Figure 3G). In addition, it showed balanced variations in long and short amplitudes, and at no time did it show similar behavior to the facial paralysis models (Figure 3H). There is a greater variation in frequency (0-50 Hz) in the movement throughout the assessment sessions (Figure 3I). This indicates that facial paralysis affects the range and frequency of whisker movement. For irreversible paralysis, the effects were permanent, and in reversible paralysis, recovery was partial, but the movement was sufficient to be compared to pre-injury states.

### Effect of facial paralysis on facial movement in mice

To assess changes in the faces of mice, a facial movement was tracked by videorecording over 2 minutes and was fragmented into frames. The difference between the first and subsequent frames was calculated by subtracting them (See materials and methods). A larger difference means that the mouse is making facial movements, whereas a smaller difference means that there is no movement. During baseline, movement can be observed in the ear, eye area, and whisker area (Figure 4-1A). For the transection injury group, movement of the ear can be observed during the 24 hours following facial paralysis. However, movement in the eye and whisker area was completely lost. By day 10 and 20, loss occurred throughout the face (Figure 4-1B). For the compression injury group, loss of movement similar to the transection group was observed on days 1 and 10. By day 20, movement can be observed in the eye, whisker, and ear areas (Figure 4-1B). Finally, the sham group never lost the ability to move its face during the days evaluated (Figure 4-1B). Therefore, facial paralysis by transection and compression affects the ability to move the mouse’s face. In contrast to previous data, transection had permanent effects, and compression had an apparent recovery.

To find out whether facial paralysis affects the mouse’s face homogeneously, facial movement analysis by areas was made: anterior (nose, lips, mouth, and whiskers), middle (cheek and eye), and posterior (ear). The frames of each video recording were obtained (n=3600) and transformed into mathematical vectors (HOGs, Figure 4A). The difference between HOGs was obtained (the subtraction between the first frame minus the subsequent ones). A greater difference indicates that the HOGs changed between frames, which translates into facial movement of the analyzed area. In the posterior area, for the three groups (transection injury, compression, and sham), a total loss of movement was not observed during all the days of evaluation, and it was comparable to the day before facial injury (Figure 4B and Figure 4-2A). In the middle and anterior area, for the transection injury group, a loss of movement was observed during the 20 days evaluated. For the compression injury group, in the middle area, there was a loss of movement in the first 8 days. Subsequently, an increase in the capacity of movement was observed until day 15, when the movement was comparable to baseline values. In the anterior area, a loss of movement was observed until day 9. From day 10, movement was recovered to baseline values (Figure 4B). In the sham group, for the middle and anterior areas, there were some days with a decrease in the movement capacity. Still, these are not significant with respect to the basal movement (Figure 4-2A). This shows that facial paralysis affects the movement of the anterior and middle areas of the face, with the eye, cheek, nose, lips, mouth, and mustache being the affected structures.

**Figure 4.**
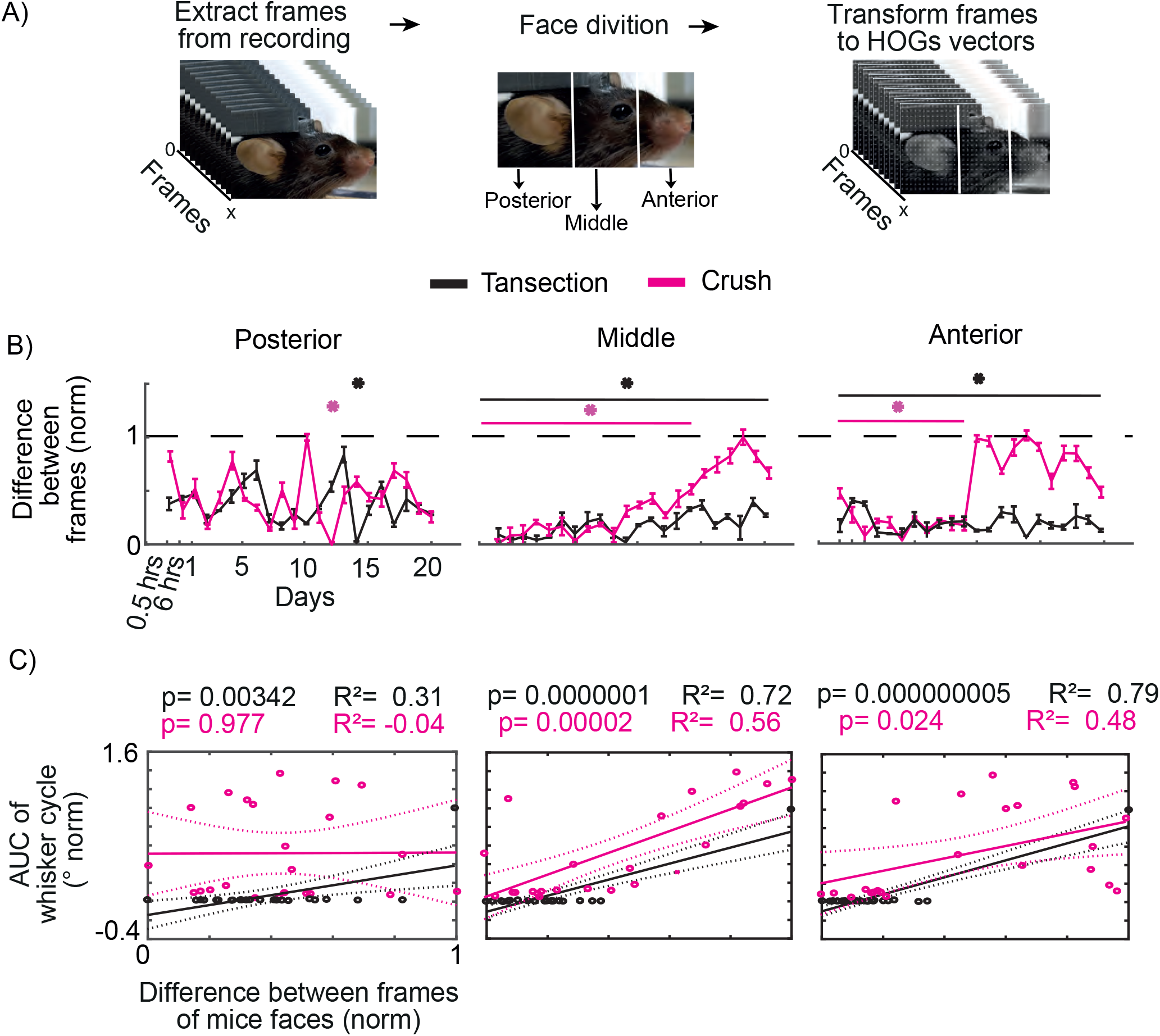
Involvement of the anterior, middle, and front parts of the face in the identification of facial paralysis. A) Schematic of the processing of the video recording of the faces of the mice. B) Differences between HOGs during 2 minutes of evaluation from half an hour to 20 days after the facial injury (n=3). The dotted line represents the baseline. The left panel symbolizes the posterior area of the face (Baseline vs Transection * p<0.05 ANOVA one way test, Tukey test p<0.05; Baseline vs Crush * p<0.05 ANOVA one way test, Tukey test p<0.05). The middle panel is the middle area of the face (Baseline vs Transection * p<0.05 ANOVA one way test, Tukey test p<0.05; Baseline vs Crush * p<0.05 ANOVA one way test, Tukey test p<0.05). The right panel shows the anterior face (Baseline vs Transection * p<0.05 ANOVA one way test, Tukey test p<0.05; Baseline vs Crush * p<0.05 ANOVA one way test, Tukey test p<0.05). C) Correlation between the difference between HOGs of the face areas of the mice (posterior, left panel; middle, middle panel and anterior, right panel, n=3) with the area under the curve of the whisker movement cycle. The points represent each of the 23 evaluations performed. The solid line shows the linear regression, and the dotted lines represent the confidence limit. Detailed statistics in Extended Data Table 4-1 and Table 4-2.

### The similarity between the evaluation of facial paralysis by means of whisker movement and the face of mice

Since facial paralysis affects the movements of the mouse’s face, the next thing we needed to know was whether the evolution of paralysis is similar in the face as in the mouse’s whisker movement. To do this, the dependence between the loss of whisker movement (range of movement determined by the area under the curve of the whisker movement cycle) and the movement of the face areas (difference in HOGs between frames) was sought. In the case of the ear area, for the transection group, a positive linear relationship is observed between the data. The opposite is true for the compression group, where the linear relationship is negative and not significant. This contrasts with the previous data, where it is observed that facial paralysis does not generate a total loss of movement of the ear area (Figure 4C). For the middle and anterior areas in both the compression and transection groups, it was observed a linearly positive relationship with the whisker movement range (Figure 4C). For sham group, the relationships of the anterior, middle, and posterior areas with the whisker movement range are linearly positive (Figure 4-2B). With this, we show that the movement of the face of mice has a positive relationship with the whisker movement, and, in the presence of facial paralysis (reversible or irreversible), the face shows a loss of movement similar to that of the whiskers. This demonstrates that video recording of the faces of mice is sufficient to detect experimental facial paralysis.

### Automatic detection of experimental facial paralysis using facial movement in mice

One of the advantages of facial paralysis is that we can detect it with the naked eye, and we can observe the moment when an apparent recovery is reached. In this sense, we now have tools that are much more precise than the human eye, such as automated image processing and computerized decision-making, which allow us to detect specific phenomena and classify them. Therefore, we seek to implement these tools to determine the presence of experimental facial paralysis. The first step was to determine a threshold for movement of the areas of the mouse face, calculated as the average of the difference in HOGs between frames throughout the days of evaluation plus the standard deviation (Figure 5A). This threshold takes into account the range of movement of all the sessions evaluated.

**Figure 5.**
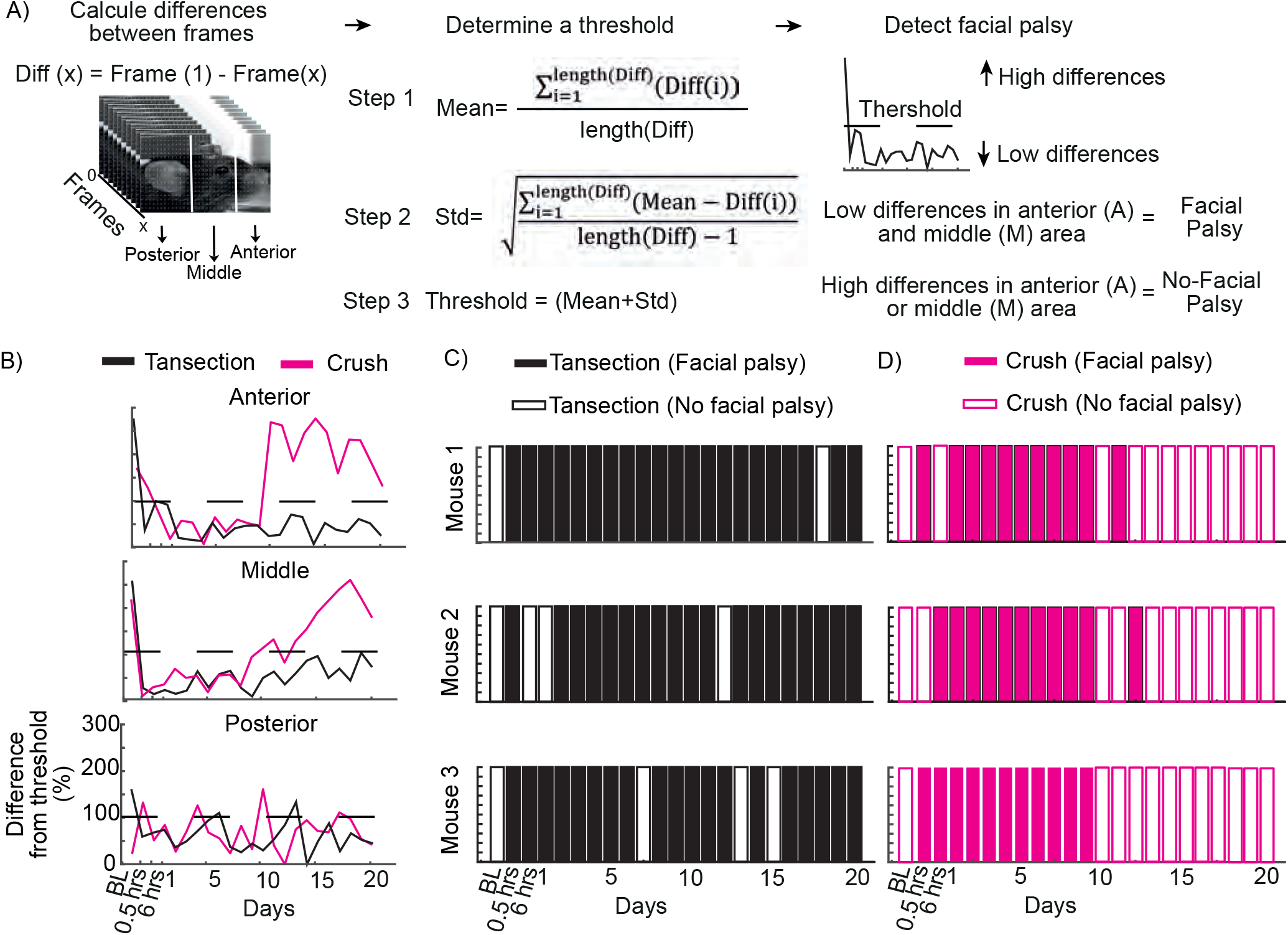
Design and implementation of the facial paralysis detection algorithm. A) Step-by-step diagram of the facial paralysis detection algorithm design. B) Percentage of change between the difference of the frames during 2 minutes of evaluation and the threshold obtained. Use of the threshold for facial paralysis detection in a binary decision-making system (facial paralysis exists or not) C) for the transection injury group (n=3) and D) for the compression injury group (n=3).

Furthermore, it was determined how different the movement of the mouse face is with respect to the threshold. For the transection group in the middle and anterior area, it can be observed below the threshold throughout the sessions evaluated (Figure 5B). In the posterior area, the motion range did not seem to remain below the threshold. For the compression injury group, in the middle area, the range of motion was below the threshold until day 13, and in the anterior area, until day 9. Afterward, the data exceed the threshold, indicating a recovery of movement in the face of the mice (Figure 5B).

To conclude that a mouse had facial paralysis, a computational decision-making model was used, which considers whether the movement in the areas of the face is below (with paralysis) or above (without paralysis) the threshold (Figure 5B, see materials and methods), which we call computational algorithm (CA). The efficiency of the CA was demonstrated by making variations in the parameters it uses, which are the areas of the face (summed or independently), data averaged by group or individually, and normalized or non-normalized data (see materials and methods). The most efficient CA is the one that takes into account the sum of the data in the anterior and middle areas, normalized and averaged by group. (Figure 5-1A; see method for more details). In this way, the CA detects facial paralysis throughout the days evaluated in mice with transection injury. For compression injury, the system detects facial paralysis within the first 12 days. Subsequently, movement is detected in the basal state (Figure 5C).

However, the AC still presents some drawbacks since the transection injury group shows days in which the system does not detect the presence of facial paralysis (Figure 5C). Therefore, the decision was made to follow up with the mice individually to find out if the facial information of each subject was sufficient to determine the presence of facial paralysis. In this sense, the number of sessions required to obtain an optimal threshold was first determined. For this purpose, the difference between the HOGs of each frame of the 23 sessions evaluated was used. These differences were averaged and used as a threshold point; 22 different thresholds were created, and in each one, a smaller number of sessions were used on average. In this way, it was observed that two sessions (a baseline evaluation and a paralyzed one) are sufficient to efficiently detect the presence of facial paralysis (Figure 5-1B). Subsequently, it was sought which session (of the 22 after facial nerve injury) was the best to create the threshold, where it is observed that no matter which session is used, the system has the same level of efficiency (Figure 5-1C). Therefore, the decision was made to use the session 24 hours after the nerve injury; the average of the movement of the anterior and middle zone was taken and taken as a threshold (Figure 6A). In this way, the CA detects facial paralysis in all mice in the days after the transection injury (Figure 6B). In the compression group, facial paralysis was detected in the first 11 days; in the case of mice one and three, there were some days in which the system detected paralysis after day 11. However, on day 17 and up to day 20, the CA no longer detected paralysis (Figure 6C). In this way, each mouse can use its facial information to track paralysis. It is capable of being used in the reversible and irreversible model and it is not necessary to use stimuli that provoke facial movement.

**Figure 6.**
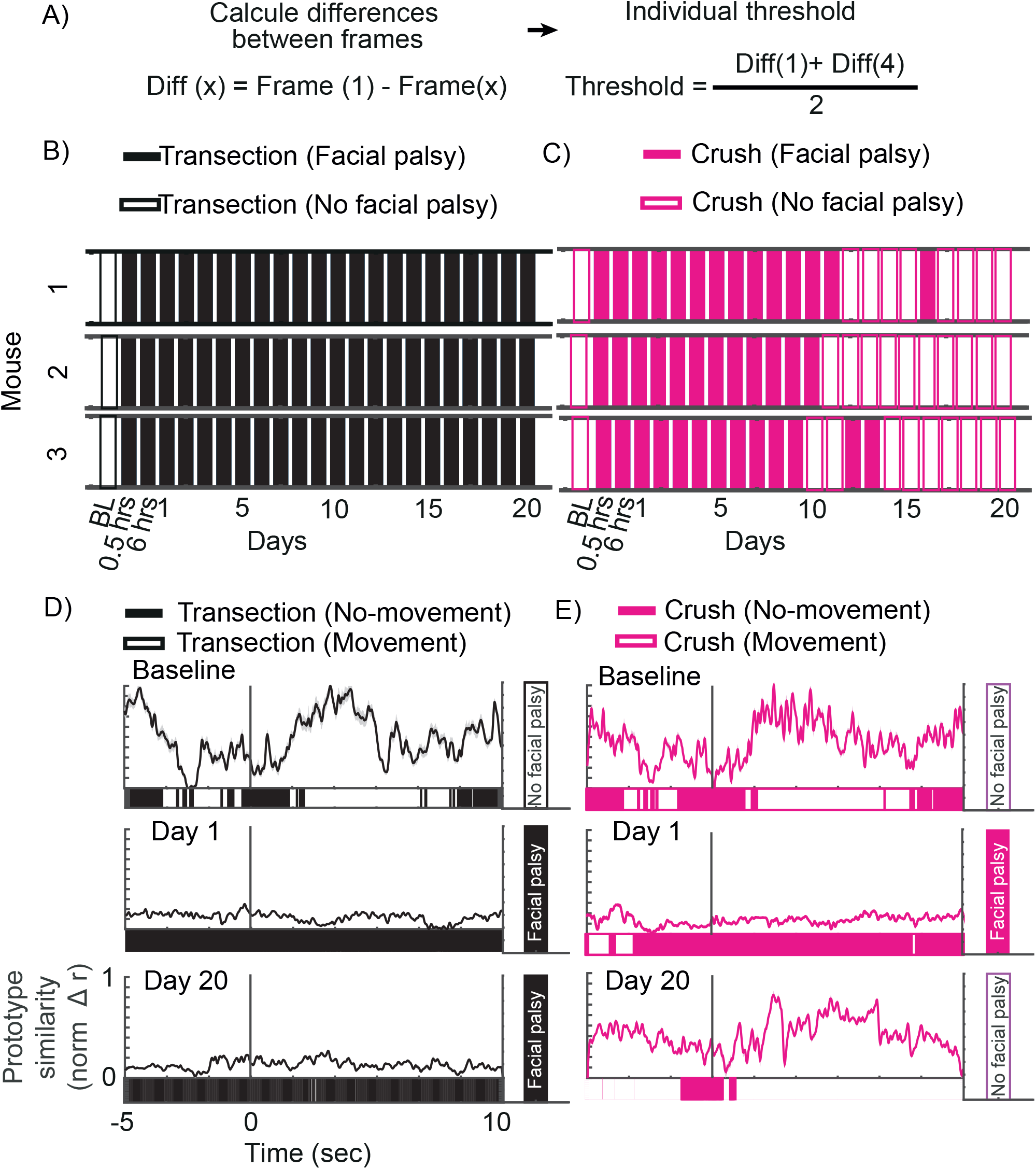
Use of the detection algorithm during the study of facial expression. A) Design of the algorithm to detect facial paralysis. Detection of facial paralysis using the computational algorithm for the transection (B, n=3) and compression (C, n=3) groups. Tracking the similarity between facial movement and the pleasurable prototype for the transection (D, n=1) and compression (E, n=1) groups. The upper panels correspond to the baseline evaluation, the middle ones to day 1, and the lower ones to day 20 after facial injury. The bar under each graph indicates the moment in which there is or is not movement. The bar on the right shows the conclusion of the computational algorithm. The black line at 0 shows the moment of sucrose release. Detailed statistics in Extended Data Table 6-2

### Detection of facial paralysis during oral stimulation with sucrose

To understand facial paralysis in depth, it is necessary to study how affected the function of facial movement is and the muscles involved. Therefore, one of the goals to be achieved is to be able to detect facial paralysis with the CA that developed during behavioral protocols. To do this, facial paralysis was detected while performing a protocol of consumption of solutions for the study of emotional facial expressions based on the protocol developed by Dolensek (2020). We released 5 drops of 4 microliters each of different solutions (bitter: quinine, sweet: sucrose, and neutral: water) to produce a facial movement associated with an emotional component (quinine-unpleasant, sucrose-pleasant and water-neutral). Such movements were video-recorded, fragmented into frames, and transformed into mathematical vectors (HOGs, Figure 6-1A).

To study facial movements associated with the consumption of a solution, the change in the face of the mice before and after stimulation was determined. For this, frames from 3 stimuli (n=210) were used, which were organized and grouped into clusters using a hierarchical tree analysis (Figure 6-1B). It can be observed that for each substance (sucrose, quinine, and water), the frames before and after stimulation are grouped into two different clusters. This indicates that the face of the mouse changes when it is orally stimulated with respect to a basal state (Figure 6-1B). Once we knew that the face changed during stimulation, we looked for the difference between the movement patterns of each administered solution. To do this, a cluster grouping was made (using a stochastic neighbor analysis of t distribution) between frames of each solution (n=120 per solution). It can be observed that each solution has its cluster (Figure 6-1C), which indicates that facial movement follows a different pattern for each solution, and this is related to the different emotions evoked by oral stimulation (Dolensek, 2020).

Regarding the above, the characteristic facial movement patterns of each solution can be extracted to generate a movement prototype (See materials and methods, Figure 6-1D). Each prototype was associated with an emotion (pleasure=sucrose, disgust=quinine, and neutral=water) and compared with each stimulus. It can be observed that stimulation with sucrose has a high similarity with the pleasant prototype but not with the neutral and disgust prototypes. Something similar occurs for quinine stimulation, which has a high similarity with the disgust prototype and not with the other two. Water follows the same pattern, where it only increases the similarity with the neutral prototype (Figure 6-1E). This shows us that facial movement is prototypical and is related to the solution administered and provokes a particular emotion.

After that, the developed CA was used to determine the presence of facial paralysis at the time of stimulation with sucrose. The system was compared with each frame during oral stimulation (n=450). In this way, we have two moments during the session, with movement and without movement. The percentage of each of the moments is obtained and the system makes its conclusion, whether there is paralysis (more than 90% of frames without movement) or not. A mouse with compression and transection injury was monitored, which, prior to nerve injury, we can observe a high similarity to the pleasure prototype after oral stimulation. In addition, during the increase in similarity, the system detects that there is facial movement. With this, the system concludes that there is no facial paralysis (Figure 6D and E). For the mouse with transection injury, we can observe on days 1 and 20 after surgery that there is no facial movement, and neither does the similarity with the pleasant prototype increase, so the system concludes that there is facial paralysis on both days (Figure 6D). For the mouse with compression injury, 24 hours after surgery, a loss of movement is observed, concluding that there is facial paralysis. However, by day 20 the movement recovers and is similar to the pleasant prototype, which results in there being no facial paralysis. In this way, the developed AC allows us to study different behaviors associated with facial movement and efficiently detects the presence of facial paralysis in the different models used.

### Facial paralysis detection during electrophysiological assessment in ALM

As a final goal, we proposed to use the designed AC during electrophysiological recording in ALM while the previously mentioned facial expression assessment protocol was being carried out (Figure 7A). Neurons associated with facial expression and movement were used. During the basal state, we can observe an increase in ALM neuronal activity at the moment that the similarity with the pleasurable prototype increases, and the algorithm detects that there is movement in the transection and compression lesion group (Figure 7B and C). Finally, the cross-covariance analysis between the firing frequency and the similarity of the facial movement with the pleasurable prototype shows that the neuronal activity occurs at the moment that the facial expression begins and is maintained during the first seconds in which it occurs. With the same analysis, comparing the firing frequency and the moments of facial movement, it is shown that the neuronal activity occurs at the moment that there is facial movement (Figure 7B and C). The cross-covariance of neuronal activity with facial expression and movement shows a significant positive correlation (Figure 7B and C).

**Figure 7.**
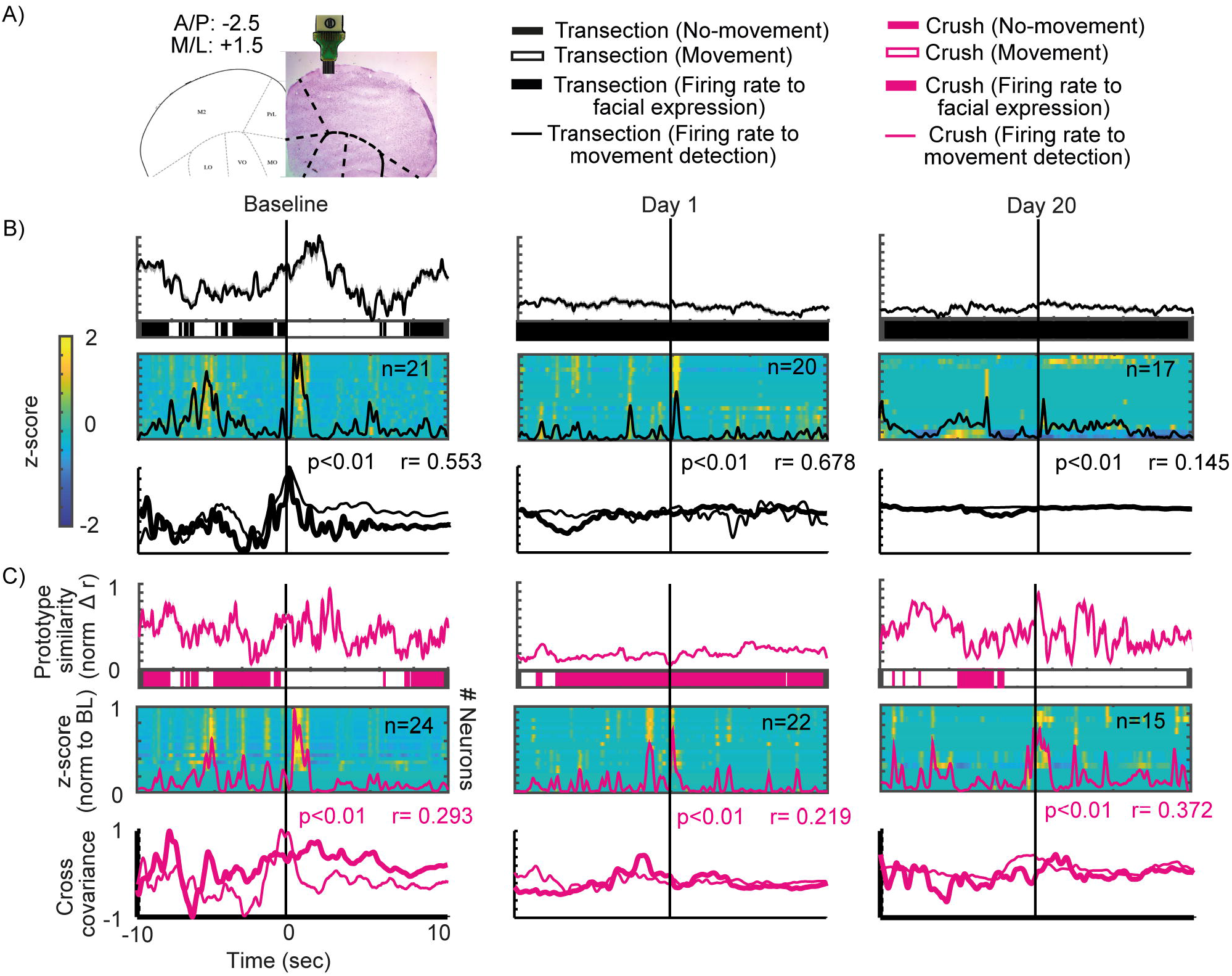
Electrophysiological recordings in ALM during facial expression and movement. A) Location of the electrode array implanted in ALM in the histological section of a recorded mouse. B and C) The top panel depicts the tracking of the similarity of the facial expression with the pleasurable prototype. In the top panel, the bar below shows the moment of facial movement. The middle panel shows the heat map of the population neuronal activity in ALM, the line inside the heat map is the population average. The bottom panel shows the cross-covariance of neuronal activity with facial expression (thick line) and movement (thin line); B) for the transection lesion group (n=2) and C) for compression (n=2). The line aligned at 0 shows the moment of facial expression onset. Detailed statistics in Extended Data Table 7-1.

For the transection injury group, 24 hours after the injury, facial expression and movement do not exist; however, an increase in the firing frequency after oral stimulation can still be seen, but it is not comparable to baseline values (Figure 7B). By day 20, facial movement does not occur, and the firing frequency decreases further compared to day 1 after the injury (Figure 7B). In this sense, the cross-covariance between the firing frequency and facial expression does not show a correlation; the same occurs between the firing frequency and the moment of movement. Likewise, both cross-covariance values show a high, significantly positive correlation (Figure 7B). This shows us that neuronal activity in ALM changes as facial paralysis progresses, and it does not occur in the same way as the loss of facial movement, which is immediate with respect to the injury.

In the compression group, 24 hours after the injury, something similar occurs as with the transection group. The firing frequency decreases with respect to basal values; there is no similarity with the pleasurable prototype, and there is no facial movement (Figure 7C). By day 20, there is an increase in the similarity to the pleasurable prototype and facial movement; likewise, the firing frequency increases to values similar to the basal state. The cross-covariance between facial expression and firing frequency shows that neuronal activity occurs after the facial expression begins. In the case of the comparison with the moment of movement, neuronal activity occurs during facial movement (Figure 7C). With this, it is shown that neuronal activity is associated with the capacity for facial movement.

To confirm that the neurons analyzed only respond to facial movement, the neuronal activity was aligned with the walking movement. It can be observed that the neurons do not show an increase in their firing frequency at the moment of initiating and maintaining walking (Figure 7-1A and B). Subsequently, the same activity was aligned with the activation of the infusion pump (see materials and methods) without administering solution; similarly to our previous result, the firing frequency has no significant modulation (Figure 7-1A and B). This demonstrates that the neuronal activity in the transection and compression group is associated with a facial response that correlates to oral stimulation.

## Discussion

In this article, we propose for the first time a model for studying facial paralysis in the mouse, which allows for easy, rapid and automated assessment of nerve injury progression. FaPDA (Facial paralysis detection algorithm) is a tool that allows us to understand the progression of experimental facial paralysis in mice, which is sensitive to this type of injury to the facial nerve, reflects changes in whisker movement, is sensitive to changes in the stereotyped movement of facial expression, and is a tool that correlates changes in neuronal activity in the premotor cortex of the rodent.

Facial nerve compression and transection techniques in murine models have been the most widely used to model facial paralysis (Chacon, Echternacht, & Leckenby, 2020) due to their similarity with the clinical prognosis. The rapid recovery that we can observe in the compression model is comparable to that of most patients with Bell’s palsy (Holland & Weiner, 2004). In contrast, the lack of recovery in transection resembles cases of poor prognosis of facial paralysis caused by herpes zoster or trauma (Holland & Weiner, 2004). However, these models have limitations since they do not capture all the variety in facial paralysis and exclude etiologies such as tumors, otitis media, or neonatal conditions. These techniques allow us to have a reversible (compression) and irreversible (transection) model capable of comparing and evaluating facial paralysis for the first time in C57BL/6 mice.

The results of our study in C57BL/6 mice corroborate previous findings in Wistar rats, showing that compression induces a rapid recovery in whisker movement amplitude. In contrast, transection shows no recovery over time (Hadlock, Kowaleski, Lo, Mackinnon, & Heaton, 2010).

In addition to assessing whisker amplitude, our study explored changes in whisker movement frequency. Jowett demonstrated that after nerve injury, rats showed a decrease in whisker spectral power (Attiah et al., 2017). Our study assessed frequency for 20 days in both models, identifying the prevalence of low frequencies (0-5 Hz) after injury. The absence of a specific stimulus to induce whisker movement limits comparability. However, our findings support the hypothesis that facial nerve injury affects not only whisker movement amplitude but also frequency.

This work proposes the study of facial paralysis using facial expression. For this purpose, we sought to use video recordings of the faces of mice without the use of any stimulus and without the need to mark or cut the whiskers (Severson et al., 2019; Vajtay et al., 2019).In addition, the importance of taking into account the complete movement of the face in the study of facial movement and conditions such as facial paralysis has been shown. (Severson et al., 2019). Our results show the loss of muscle function in two of the three facial regions studied (middle and anterior area). It could be related to the impact of the facial injury on the musculature of the mouse ear and its need for sound sensation. (Cakmak, 2019; Clayton et al., 2024).

The effect of facial paralysis on whisker movement is related to the mechanical technique used to produce the injury (Hadlock et al., 2010).

In contrast, the results show a loss of facial movement function temporally similar to the loss of whisker movement. They are also dependent on the type of injury to the facial nerve. This is observed in the anterior and middle areas of the face.

Currently, the use of digital tools and artificial intelligence for the study of the complexity of facial movement has been shown more frequently (Syeda et al., 2024; Tanaka et al., 2023).

In order to automatically detect facial paralysis, this work proposes a bimodal detection algorithm, obtaining information on the movement of the mouse’s face in three segments since there is a high participation of its structures in facial movement. (Le Moene & Larsson, 2023; Tanaka et al., 2023). The results show us the importance of using the middle and anterior area of the mouse’s face for the automated detection of facial paralysis, despite the fact that the ear area is essential for generating a facial expression (Dolensek et al., 2020).

Furthermore, it is shown that the information specific to each mouse is sufficient to feed the algorithm and monitor the facial paralysis of each subject. It can help us understand whether this condition has unique effects on each mouse or can be studied further as a group.

In this study, we propose the use of artificial vision as a method to characterize the effects of facial nerve injury on musculature movements. For this purpose, we used the facial response elicited by a sucrose solution, as it has been shown that the facial changes are proportional to the intensity of the hedonic stimuli (Le Moene & Larsson, 2023) and remain consistent when evaluated in a head-fixed model through video recording analysis, allowing the determination of the internal state of mice in a quantitative manner (W. R. Li et al., 2023). As such, the reduced motion observed after the facial nerve injury seems to be related to the inability to perform the facial movements rather than a devaluation of the reward being delivered.

To corroborate the usefulness of this tool in the assessment of facial paralysis in mice, we analyzed its neural correlates in the anterolateral motor cortex, a possible homolog of primates’ premotor cortex. Our data confirm that spontaneous orofacial responses are initiated and executed in this cortex (Giordano et al., 2023) and that the neural modulation associated with orofacial movement disappears under conditions of experimental facial paralysis irreversible.

On the contrary, our data also suggest that neuronal modulation of the premotor cortex is reestablished on day 20 after nerve injury in the reversible model. Therefore, the experimental model works to see fine changes at the cortical level associated with the natural history of experimental facial paralysis.

Further in-depth experiments are still required to analyze aspects of planning and execution, also attributed to the anterolateral motor cortex (N. Li et al., 2015), and that in other experimental conditions give us the perspective of treatment with neuroprostheses or specific neuromodulation applied to experimental facial paralysis in rodents.

---

Figure 2-1.

**Detection of the onset moment of whisker movement**. A) Tracking of a whisker movement over 4 seconds. The green line shows the moments of greatest change in the signal. B) Detection of the high points of the signal (peaks) marked by the blue circles; the green line represents the moments of abrupt changes along the signal, and the black point shows the point with the greatest change (change point). C) Amplitudes formed in the signal before and after the change point for the transection group, in the upper panel (t-test * p < 0.05); compression, in the middle panel (t-test * p < 0.05) and sham, in the lower panel (t-test * p < 0.05). Detailed statistics in Extended Data Table 2-1

Figure 2-2.

**The cycle of whisker movement contralateral to facial injury**. Whisker movement was monitored in one mouse per group, (A) transection and (B) crush, prior to surgery and on days 1, 10, and 20 postoperatively. The graphs on the left of each column represent the tracking of whisker movement dynamics over 2 seconds of assessment, and the movement cycle is shown on the right.

Figure 4-1.

**Facial movement between frames during facial video recording of mice**. A) Heat map of a representative mouse before and B) after facial nerve injury (days 1, 10, and 20); a representative mouse for the transection group in the upper panels; crush in the middle panels; and sham in the lower panels.

Figure 4-2.

**Assessment of mouse facial areas in the sham group**. A) Differences between HOGs during 2 minutes of assessment (n=3), from half an hour to 20 days after a facial injury. The dotted line represents the baseline. The left panel symbolizes the posterior area of the face (Baseline vs Sham * p>0.05 ANOVA one-way test). The middle panel is the middle area of the face (Baseline vs Sham * p>0.05 ANOVA one-way test). The right panel is the anterior area of the face (Baseline vs Sham * p>0.05 ANOVA one-way test). Correlation between the difference between the HOGs of the facial areas of the mice (posterior, left panel; middle, middle panel and anterior, right panel, n=3) and the area under the whisker movement cycle curve. The dots represent each of the 23 assessments performed. The solid line shows the linear regression, and the dotted lines represent the confidence limit. Detailed statistics in Extended Data Table 4-3.

Figure 5-1.

**Facial paralysis detection algorithm efficiency**. A) Level of efficiency of the different algorithms designed to detect facial paralysis in continuous and random data. The rectangle indicates the designed algorithm with the highest efficiency in detecting paralysis. B) Efficiency of the algorithm using different amounts of post-facial injury assessments averaged together; each bar indicates the number of assessments used. Efficiency of the algorithm when averaging the baseline with one post-facial injury assessment; each bar indicates the assessment used with the baseline.

Figure 6-1.

**Facial expression detection with oral stimulation of different solutions**. A) Schematic of the analysis process of the video recordings to detect facial expressions. B) Heat map of the correlation of the frames of 3 stimuli (n=210 in one mouse) before and after the release of sucrose, water, and quinine, sorted by clusters. C) Grouping by clusters of the frames after oral stimulation with sucrose, water, and quinine. D) Schematic of the creation of the pleasant, unpleasant, and neutral prototypes. E) Similarity of the prototypes with the moments of oral stimulation with the different solutions (sucrose, quinine and water) in three mice. The dotted line at 0 shows the moment of the release of solutions. Detailed statistics in Extended Data Table 6-1.

Figure 7-1.

**Electrophysiological recordings in ALM during walking and activation of the solution infusion pump**. The upper panel shows the heat map of population activity in ALM associated with the activation of the solution infusion pump, and the lower panel shows mouse walking for A) the transection group (n=2) and B) compression (n=2). The black line aligned at 0 indicates the moment when the pump is activated and walking begins. Detailed statistics in Extended Data Table 7-2.

Table 1-1

**Statistical details in the forces applied in facial nerve**. Difference between the forces applied in facial nerve in the crush group (mouse 1 vs. mouse 2 vs mouse 3). Significance level =0.05.

Table 2-1

**Statistical details in whisker movement**. Difference between the amplitudes before and after the change point in transection, crush, and sham groups. Significance level =0.05.

Table 3-1

**Statistical details in whisker movement with facial paralysis**. Difference in area under the curve between baseline vs. days post facial paralysis in transection, crush, and sham groups. Significance level =0.05.

Table 3-2

**Statistical details in the proportion of high and low amplitudes in transection group**. Difference between the baseline day vs. days post facial paralysis. Significance level =0.05.

Table 3-3

**Statistical details in the proportion of high and low amplitudes in crush group**. Difference between the baseline day vs. days post facial paralysis. Significance level =0.05.

Table 3-4

**Statistical details in the proportion of high and low amplitudes in sham group**. Difference between the baseline day vs. days post facial paralysis. Significance level =0.05.

Table 4-1

**Statistical details in the differences between frames in transection group**. Difference between the first frame with the others in the video, comparison between baseline vs days post facial paralysis. Significance level =0.05.

Table 4-2

**Statistical details in the differences between frames in crush group**. Difference between the first frame with the others in the video, comparison between baseline vs days post facial paralysis. Significance level =0.05.

Table 4-3

**Statistical details in the differences between frames in sham group**. Difference between the first frame with the others in the video, comparison between baseline vs days post facial paralysis. Significance level =0.05.

Table 6-1

**Statistical details in facial expression before and after oral stimulation with solutions**. Comparison between 10 seconds pre and post-stimulation with sucrose, quinine, or water using the pleasure, disgust, and neutral prototype. Significance level =0.05.

Table 6-2

**Statistical details in facial expression between baseline vs facial paralysis**. Similarity of pleasure prototype for 10 seconds after oral stimulation with sucrose between baseline, day 1, and day 20 post facial paralysis in transection and crush group. Significance level =0.05.

Table 7-1

**Statistical details in population neuronal activity in ALM pre and post-facial expression**. Differences between z-score pre and post-oral stimulation with sucrose in population neuronal activity of two mice. Significance level =0.05.

Table 7-2

**Statistical details in population neuronal activity in ALM pre and post-control events**. Differences between z-score pre and post-walking and activation of a stepper motor in population neuronal activity of two mice. Significance level =0.05.

**Extended Data 1**

Extended data includes a copy of the GitHub repository. Contains a copy of code with two sections. Section 1: code to develop FaPDA with any video recording information of facial paralysis. Section 2: code to use FaPDA in video recordings of mice with facial paralysis.

## Notes

### Competing Interest Statement

The authors have declared no competing interest.

## References

Attiah, M. A., de Vries, J., Richardson, A. G., & Lucas, T. H. (2017). A Rodent Model of Dynamic Facial Reanimation Using Functional Electrical Stimulation. Front Neurosci, 11, 193. doi:10.3389/fnins.2017.00193

Cakmak, Y. O. (2019). Concerning Auricular Vagal Nerve Stimulation: Occult Neural Networks. Front Hum Neurosci, 13, 421. doi:10.3389/fnhum.2019.00421

Chacon, M. A., Echternacht, S. R., & Leckenby, J. I. (2020). Outcome measures of facial nerve regeneration: A review of murine model systems. Ann Anat, 227, 151410. doi:10.1016/j.aanat.2019.07.011

Clayton, K. K., Stecyk, K. S., Guo, A. A., Chambers, A. R., Chen, K., Hancock, K. E., & Polley, D. B. (2024). Sound elicits stereotyped facial movements that provide a sensitive index of hearing abilities in mice. Current Biology, 34(8), 1605-1620.e1605. doi:10.1016/j.cub.2024.02.057

Dolensek, N., Gehrlach, D. A., Klein, A. S., & Gogolla, N. (2020). Facial expressions of emotion states and their neuronal correlates in mice. Science, 368(6486), 89–94. doi:doi:10.1126/science.aaz9468

Giordano, N., Alia, C., Fruzzetti, L., Pasquini, M., Palla, G., Mazzoni, A., … Caleo, M. (2023). Fast-Spiking Interneurons of the Premotor Cortex Contribute to Initiation and Execution of Spontaneous Actions. J Neurosci, 43(23), 4234–4250. doi:10.1523/JNEUROSCI.0750-22.2023

Hadlock, T. A., Kowaleski, J., Lo, D., Mackinnon, S. E., & Heaton, J. T. (2010). Rodent facial nerve recovery after selected lesions and repair techniques. Plast Reconstr Surg, 125(1), 99–109. doi:10.1097/PRS.0b013e3181c2a5ea

Haiss, F., & Schwarz, C. (2005). Spatial segregation of different modes of movement control in the whisker representation of rat primary motor cortex. J Neurosci, 25(6), 1579–1587. doi:10.1523/JNEUROSCI.3760-04.2005

Holland, N. J., & Weiner, G. M. (2004). Recent developments in Bell’s palsy. BMJ, 329(7465), 553–557. doi:10.1136/bmj.329.7465.553

Klingner, C. M., Volk, G. F., Maertin, A., Brodoehl, S., Burmeister, H. P., Guntinas-Lichius, O., & Witte, O. W. (2011). Cortical reorganization in Bell’s palsy. Restor Neurol Neurosci, 29(3), 203–214. doi:10.3233/RNN-2011-0592

Le Moene, O., & Larsson, M. (2023). A New Tool for Quantifying Mouse Facial Expressions. eNeuro, 10(2). doi:10.1523/ENEURO.0349-22.2022

Li, N., Chen, T. W., Guo, Z. V., Gerfen, C. R., & Svoboda, K. (2015). A motor cortex circuit for motor planning and movement. Nature, 519(7541), 51–56. doi:10.1038/nature14178

Li, W. R., Nakano, T., Mizutani, K., Matsubara, T., Kawatani, M., Mukai, Y., … Yamashita, T. (2023). Neural mechanisms underlying uninstructed orofacial movements during reward-based learning behaviors. Curr Biol, 33(16), 3436–3451 e3437. doi:10.1016/j.cub.2023.07.013

Maspero, C., Farronato, M., Guenza, G., & Farronato, D. (2017). Long term results of idiopathic hemifacial palsy: Orthodontic and surgical multidisciplinary management. Oral and Maxillofacial Surgery Cases, 3(4), 86–101. doi:10.1016/j.omsc.2017.08.002

Munera, A., Cuestas, D. M., & Troncoso, J. (2012). Peripheral facial nerve lesions induce changes in the firing properties of primary motor cortex layer 5 pyramidal cells. Neuroscience, 223, 140–151. doi:10.1016/j.neuroscience.2012.07.063

Severson, K. S., Xu, D., Yang, H., & O’Connor, D. H. (2019). Coding of whisker motion across the mouse face. Elife, 8. doi:10.7554/eLife.41535

Song, W., Dai, M., Xuan, L., Cao, Z., Zhou, S., Lang, C., … Kong, J. (2017). Sensorimotor Cortical Neuroplasticity in the Early Stage of Bell’s Palsy. Neural Plast, 2017, 8796239. doi:10.1155/2017/8796239

Sugita, T., Murakami, S., Yanagihara, N., Fujiwara, Y., Hirata, Y., & Kurata, T. (1995). Facial nerve paralysis induced by herpes simplex virus in mice: an animal model of acute and transient facial paralysis. Ann Otol Rhinol Laryngol, 104(7), 574–581. doi:10.1177/000348949510400713

Syeda, A., Zhong, L., Tung, R., Long, W., Pachitariu, M., & Stringer, C. (2024). Facemap: a framework for modeling neural activity based on orofacial tracking. Nat Neurosci, 27(1), 187–195. doi:10.1038/s41593-023-01490-6

Tanaka, Y., Nakata, T., Hibino, H., Nishiyama, M., & Ino, D. (2023). Classification of multiple emotional states from facial expressions in head-fixed mice using a deep learning-based image analysis. PLoS One, 18(7), e0288930. doi:10.1371/journal.pone.0288930

Urrego, D., Munera, A., & Troncoso, J. (2011). [Peripheral facial nerve lesion induced long-term dendritic retraction in pyramidal cortico-facial neurons]. Biomedica, 31(4), 560–569. doi:10.1590/S0120-41572011000400011

Vajtay, T. J., Bandi, A., Upadhyay, A., Swerdel, M. R., Hart, R. P., Lee, C. R., & Margolis, D. J. (2019). Optogenetic and transcriptomic interrogation of enhanced muscle function in the paralyzed mouse whisker pad. J Neurophysiol, 121(4), 1491–1500. doi:10.1152/jn.00837.2018

Walker, N. R., Mistry, R. K., & Mazzoni, T. (2024). Facial Nerve Palsy. In StatPearls. Treasure Island (FL) ineligible companies. Disclosure: Rakesh Mistry declares no relevant financial relationships with ineligible companies. Disclosure: Thomas Mazzoni declares no relevant financial relationships with ineligible companies.

Zeng, Z., Bai, Y., Hao, W., Zhang, T., Yang, J., Wu, F., & Li, X. (2024). Elevated TRPV2 expression in the facial nerve of rats by cold stimulation: Implications for Bell’s palsy. J Stomatol Oral Maxillofac Surg, 101895. doi:10.1016/j.jormas.2024.101895

