## supplemental for "FaPDA: Facial paralysis detection algorithm applied in mice"

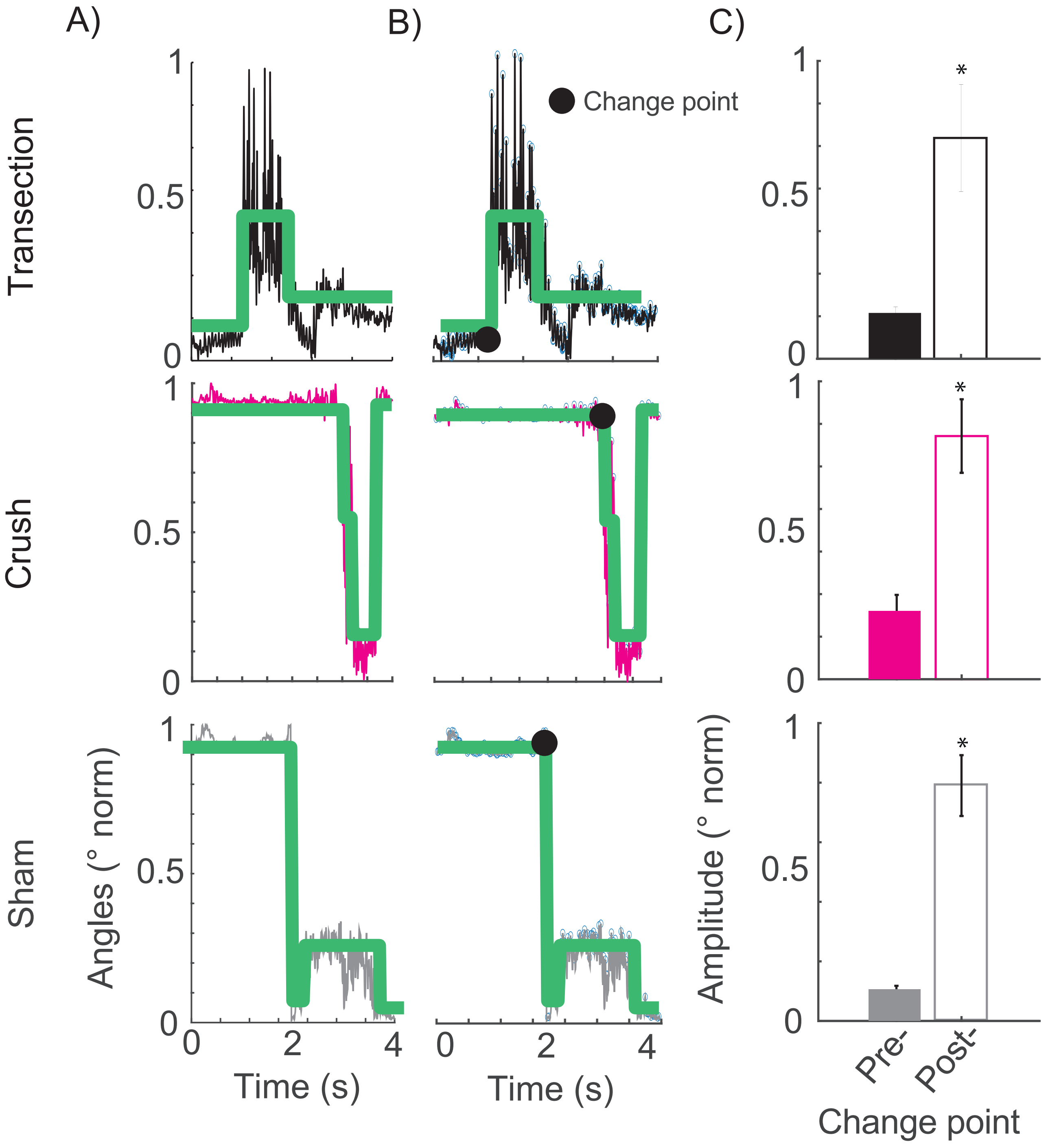

Figure 2-1.
Detection of the onset moment of whisker movement. A) Tracking of a whisker movement over 4 seconds. The green line shows the moments of greatest change in the signal. B) Detection of the high points of the signal (peaks) marked by the blue circles; the green line represents the moments of abrupt changes along the signal, and the black point shows the point with the greatest change (change point). C) Amplitudes formed in the signal before and after the change point for the transection group, in the upper panel (t-test * $p < 0.05$); compression, in the middle panel (t-test * $p < 0.05$) and sham, in the lower panel (t-test * $p < 0.05$). Detailed statistics in Extended Data Table 2-1

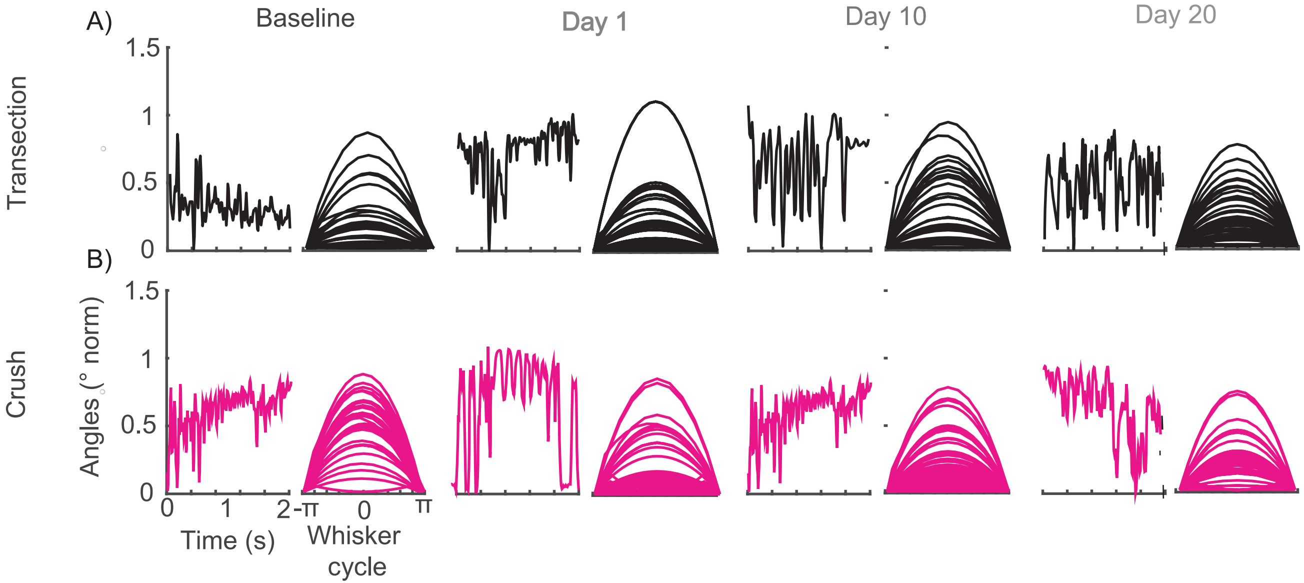

Figure 2-2.
The cycle of whisker movement contralateral to facial injury. Whisker movement was monitored in one mouse per group, (A) transection and (B) crush, prior to surgery and on days 1, 10, and 20 postoperatively. The graphs on the left of each column represent the tracking of whisker movement dynamics over 2 seconds of assessment, and the movement cycle is shown on the right.

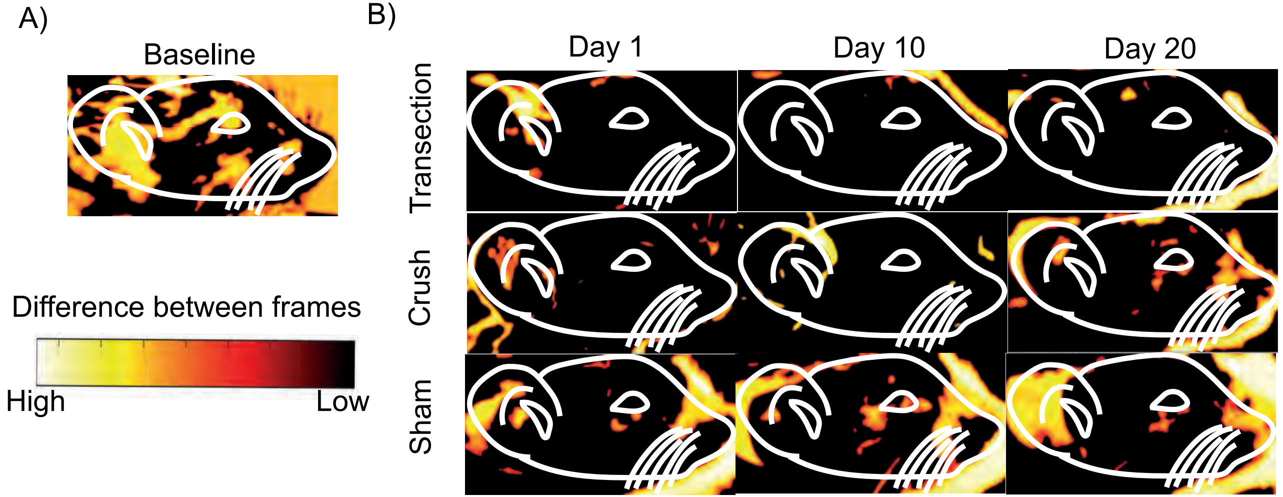

Figure 4-1.
Facial movement between frames during facial video recording of mice. A) Heat map of a representative mouse before and B) after facial nerve injury (days 1, 10, and 20); a representative mouse for the transection group in the upper panels; crush in the middle panels; and sham in the lower panels.

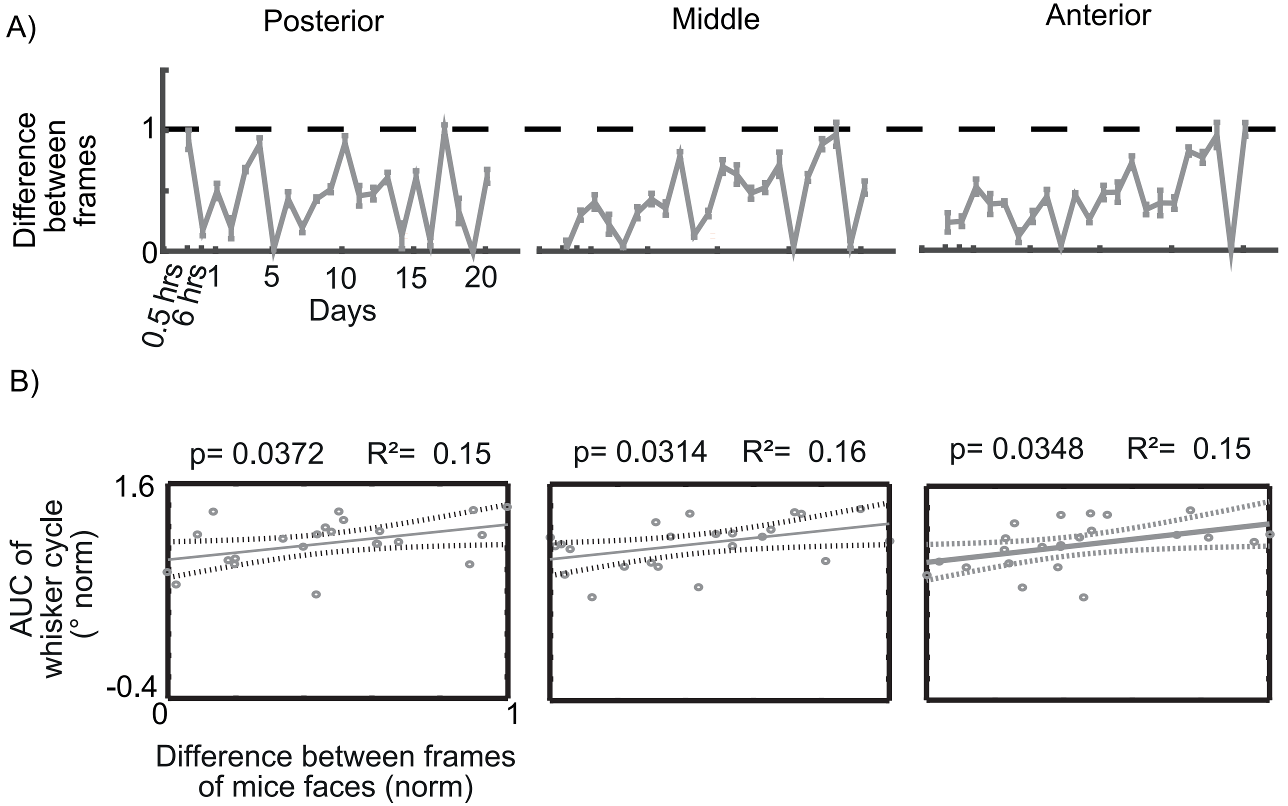

Figure 4-2.

Assessment of mouse facial areas in the sham group. A) Differences between HOGs during 2 minutes of assessment (n=3), from half an hour to 20 days after a facial injury. The dotted line represents the baseline. The left panel symbolizes the posterior area of the face (Baseline vs Sham * $p>0.05$ ANOVA one-way test). The middle panel is the middle area of the face (Baseline vs Sham * $p>0.05$ ANOVA one-way test). The right panel is the anterior area of the face (Baseline vs Sham * $p>0.05$ ANOVA one-way test). B) Correlation between the difference between the HOGs of the facial areas of the mice (posterior, left panel; middle, middle panel and anterior, right panel, n=3) and the area under the whisker movement cycle curve. The dots represent each of the 23 assessments performed. The solid line shows the linear regression, and the dotted lines represent the confidence limit. Detailed statistics in Extended Data Table 4-3.

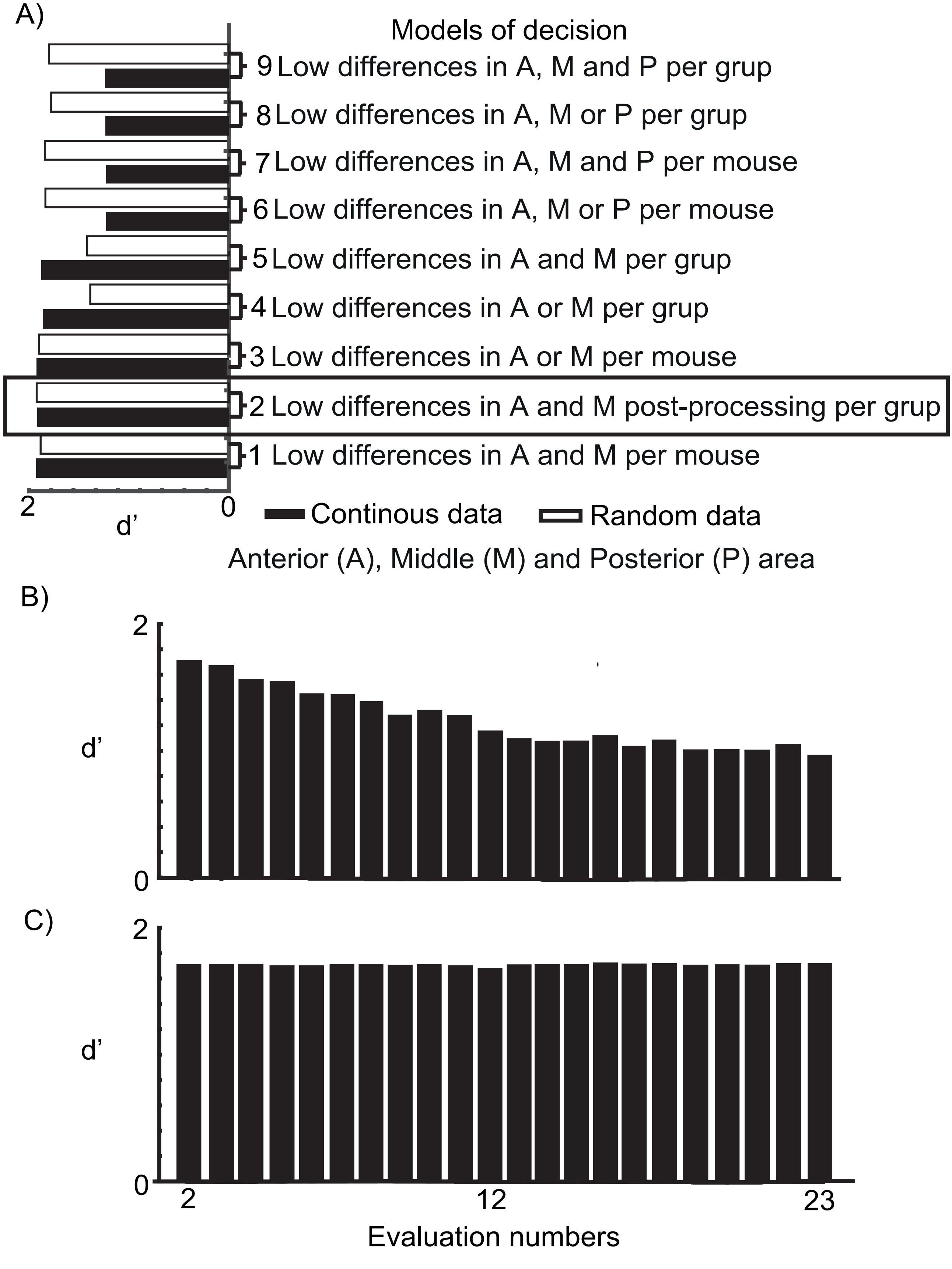

Figure 5-1.
Facial paralysis detection algorithm efficiency. A) Level of efficiency of the different algorithms designed to detect facial paralysis in continuous and random data. The rectangle indicates the designed algorithm with the highest efficiency in detecting paralysis. B) Efficiency of the algorithm using different amounts of post-facial injury assessments averaged together; each bar indicates the number of assessments used. C) Efficiency of the algorithm when averaging the baseline with one post-facial injury assessment; each bar indicates the assessment used with the baseline.

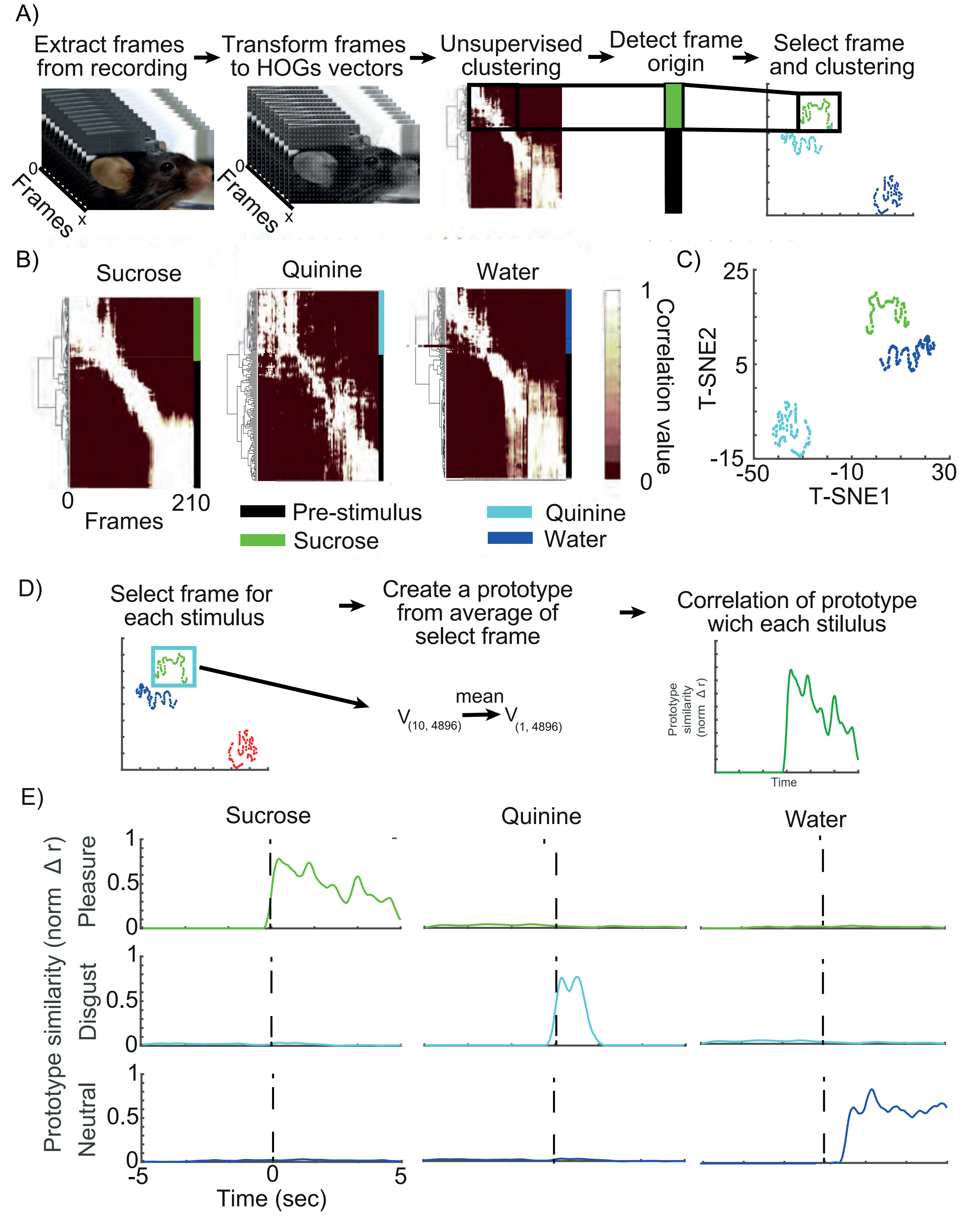

Figure 6-1.

Facial expression detection with oral stimulation of different solutions. A) Schematic of the analysis process of the video recordings to detect facial expressions. B) Heat map of the correlation of the frames of 3 stimuli (n=210 in one mouse) before and after the release of sucrose, water, and quinine, sorted by clusters. C) Grouping by clusters of the frames after oral stimulation with sucrose, water, and quinine. D) Schematic of the creation of the pleasant, unpleasant, and neutral prototypes. E) Similarity of the prototypes with the moments of oral stimulation with the different solutions (sucrose, quinine and water) in three mice. The dotted line at 0 shows the moment of the release of solutions. Detailed statistics in Extended Data Table 6-1.

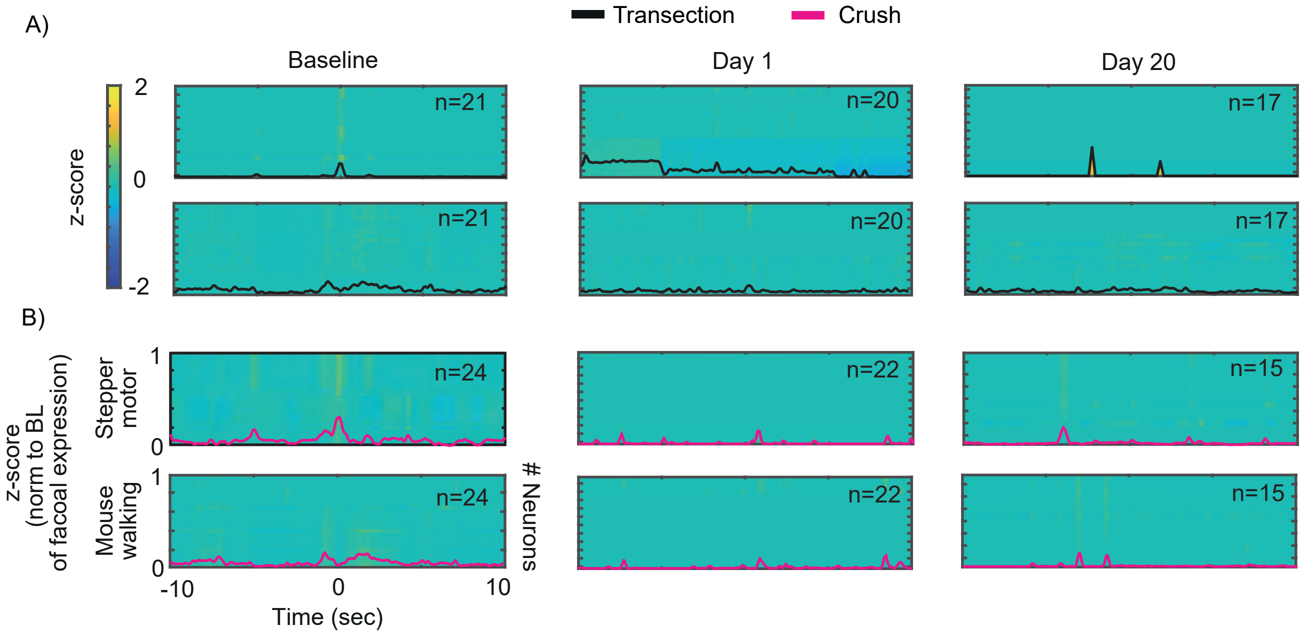

Figure 7-1.
Electrophysiological recordings in ALM during walking and activation of the solution infusion pump. The upper panel shows the heat map of population activity in ALM associated with the activation of the solution infusion pump, and the lower panel shows mouse walking for A) the transection group (n=2) and B) compression (n=2). The black line aligned at 0 indicates the moment when the pump is activated and walking begins. Detailed statistics in Extended Data Table 7-2.

**Table 1-1**

| one way ANOVA | | |
|---|---|---|
| df | F value | p value |
| 2 | 0.1081 | 0.8975 |

**Statistical details in the forces applied in facial nerve.** Difference between the forces applied in facial nerve in the crush group (mouse 1 vs mouse 2 vs mouse 3). Significance level =0.05.

**Table 2-1**

| T-test | | | |
|---|---|---|---|
| Facial palsy model | df | sd value | p value |
| Transection | 24 | 9.128 | 0.0037 |
| Crush | 14 | 18.309 | 0.00067 |
| Sham | 28 | 13.83 | 0.00000031 |

**Statistical details in whisker movement.** Difference between the amplitudes before and after the change point in transection, crush and sham groups. Significance level =0.05.

**Table 3-1**

| Transection | | | | | Crush | | | | Sham | | |
|---|---|---|---|---|---|---|---|---|---|---|---|
| Analysis: one way ANOVA | | | | | | | | | | | |
| df | F value | p value | | | df | F vaue | p value | | df | F vaue | p value |
| 22 | 561.408 | 1E-12 | | | 22 | 10.9544 | 1.41E-36 | | 22 | 10.1836569 | 6.73E-34 |
| Post hoc Tukye | | | | | Post hoc Tukye | | | | Post hoc Tukye | | |
| Comparationlow confidence interval | low confidence inteval | high confidence interval | p value | | low confidence inteval | high confidence interval | p value | | low confidence inteval | high confidence interval | p value |
| .5 hrs | 0.7138417 | 0.78100716 | 1E-27 | | 0.10539581 | 1.01729824 | 0.00204952 | | -0.1014197 | 0.09627043 | 0.99999871 |
| 6 hrs | 0.70963214 | 0.7767976 | 1E-27 | | 0.11144431 | 1.1621428 | 0.00257567 | | -0.0419804 | 0.15570974 | 0.91080611 |
| Day 1 | 0.71776382 | 0.78492928 | 1E-27 | | 0.11155378 | 1.16225227 | 0.00256718 | | -0.1939849 | 0.00370524 | 0.07719326 |
| Day 2 | 0.71317778 | 0.78034324 | 1E-27 | | 0.09462982 | 1.1453283 | 0.00423982 | | -0.0615361 | 0.13615399 | 0.9995366 |
| Day 3 | 0.71149686 | 0.77866232 | 1E-27 | | 0.10539581 | 1.1560943 | 0.00308746 | | -0.11953 | 0.07816014 | 0.9999987 |
| Day 4 | 0.71405474 | 0.7812202 | 1E-27 | | 0.11896914 | 1.16966763 | 0.00204952 | | -0.0573448 | 0.14034532 | 0.99776969 |
| Day 5 | 0.70969296 | 0.77685841 | 1E-27 | | 0.13520273 | 1.18590121 | 0.00123783 | | -0.009691 | 0.18799911 | 0.14591316 |
| Day 6 | 0.71384061 | 0.78100607 | 1E-27 | | 0.14004315 | 1.19074164 | 0.0010619 | | -0.1348413 | 0.06284883 | 0.9997347 |
| Day 7 | 0.7133756 | 0.78054105 | 1E-27 | | 0.10468558 | 1.15538407 | 0.00315342 | | -0.0719621 | 0.12572799 | 0.99999697 |
| Day 8 | 0.71431865 | 0.78148411 | 1E-27 | | 0.11041299 | 1.16111147 | 0.00265692 | | 0.02053692 | 0.21822701 | 0.00275958 |
| Day 9 | 0.71497784 | 0.7821433 | 1E-27 | | 0.079028 | 1.12972649 | 0.00663004 | | -0.1715719 | 0.02611825 | 0.52688087 |

| | | | | | | | | | | | |
|---|---|---|---|---|---|---|---|---|---|---|---|
| Day 10 | 0.71354805 | 0.78071351 | 1E-27 | | 0.09086726 | 1.14156575 | 0.00472899 | | -0.1971231 | 0.00056703 | 0.05353686 |
| Day 11 | 0.71086172 | 0.77802718 | 1E-27 | | 0.04650857 | 1.09720706 | 0.01602326 | | -0.1556569 | 0.04203319 | 0.91155529 |
| Day 12 | 0.70782801 | 0.77499347 | 1E-27 | | -0.104445 | 0.94625349 | 0.34646532 | | -0.1414506 | 0.0562395 | 0.99678469 |
| Day 13 | 0.71264314 | 0.7798086 | 1E-27 | | -0.0737284 | 0.97697007 | 0.21636959 | | -0.1104895 | 0.08720056 | 0.99999871 |
| Day 14 | 0.71089786 | 0.77806332 | 1E-27 | | -0.4665345 | 0.58416403 | 0.99999871 | | -0.1935167 | 0.00417336 | 0.08137375 |
| Day 15 | 0.71188235 | 0.77904781 | 1E-27 | | -0.2483285 | 0.80237 | 0.96215568 | | -0.1140049 | 0.08368516 | 0.99999871 |
| Day 16 | 0.71241675 | 0.7795822 | 1E-27 | | -0.7951749 | 0.25552358 | 0.97154189 | | -0.1339803 | 0.06370977 | 0.9998193 |
| Day 17 | 0.71431894 | 0.7814844 | 1E-27 | | -0.7072401 | 0.34345839 | 0.99988163 | | -0.2049204 | -0.0072303 | 0.01973809 |
| Day 18 | 0.71193691 | 0.77910237 | 1E-27 | | -0.7388313 | 0.31186723 | 0.99860211 | | -0.1247464 | 0.07294374 | 0.99999784 |
| Day 19 | 0.71266442 | 0.77982988 | 1E-27 | | -0.5595911 | 0.49110738 | 0.99999871 | | -0.0367969 | 0.16089321 | 0.81663512 |
| Day 20 | 0.71335228 | 0.78051774 | 1E-27 | | -0.6508563 | 0.39984219 | 0.99999856 | | -0.1424711 | 0.05521902 | 0.99556646 |

**Statistical details in whisker movement with facial parlysis.** Difference in area under the curve between baseline vs days post facial paralysis in transection, crush and sham groups. Significance level =0.05.

**Table 3-2**

| Transection | | | | | | | |
|---|---|---|---|---|---|---|---|
| Chi- square test | | | | | | | |
| | High amplitudes | | | | Low amplitudes | | |
| | p value | percentage | N | | p value | percentage | N |
| .5 hrs | 6.33E-15 | 0.0443066 | 1 | | 4.11E-14 | 99.9556934 | 2256 |
| 6 hrs | 6.66E-15 | 0.10638298 | 1 | | 4.36E-14 | 99.893617 | 939 |
| Day 1 | 9.88E-15 | 0.4950495 | 1 | | 6.28E-14 | 99.5049505 | 201 |
| Day 2 | 9.88E-15 | 0.4950495 | 1 | | 6.28E-14 | 99.5049505 | 201 |
| Day 3 | 9.88E-15 | 0.4950495 | 1 | | 6.28E-14 | 99.5049505 | 201 |
| Day 4 | 9.88E-15 | 0.4950495 | 1 | | 6.28E-14 | 99.5049505 | 201 |
| Day 5 | 9.88E-15 | 0.4950495 | 1 | | 6.28E-14 | 99.5049505 | 201 |
| Day 6 | 9.88E-15 | 0.4950495 | 1 | | 6.28E-14 | 99.5049505 | 201 |
| Day 7 | 9.88E-15 | 0.4950495 | 1 | | 6.28E-14 | 99.5049505 | 201 |
| Day 8 | 9.88E-15 | 0.4950495 | 1 | | 6.28E-14 | 99.5049505 | 201 |
| Day 9 | 9.88E-15 | 0.4950495 | 1 | | 6.28E-14 | 99.5049505 | 201 |
| Day 10 | 9.88E-15 | 0.4950495 | 1 | | 6.28E-14 | 99.5049505 | 201 |
| Day 11 | 9.88E-15 | 0.4950495 | 1 | | 6.28E-14 | 99.5049505 | 201 |
| Day 12 | 9.88E-15 | 0.4950495 | 1 | | 6.28E-14 | 99.5049505 | 201 |
| Day 13 | 9.88E-15 | 0.4950495 | 1 | | 6.28E-14 | 99.5049505 | 201 |
| Day 14 | 9.88E-15 | 0.4950495 | 1 | | 6.28E-14 | 99.5049505 | 201 |
| Day 15 | 9.88E-15 | 0.4950495 | 1 | | 6.28E-14 | 99.5049505 | 201 |
| Day 16 | 9.88E-15 | 0.4950495 | 1 | | 6.28E-14 | 99.5049505 | 201 |
| Day 17 | 9.10E-15 | 0.4048583 | 1 | | 5.77E-14 | 99.5951417 | 246 |
| Day 18 | 9.10E-15 | 0.4048583 | 1 | | 5.77E-14 | 99.5951417 | 246 |
| Day 19 | 9.10E-15 | 0.4048583 | 1 | | 5.77E-14 | 99.5951417 | 246 |
| Day 20 | 9.10E-15 | 0.4048583 | 1 | | 5.77E-14 | 99.5951417 | 246 |

**Statistical details in the proportion of high and low amplitudes in transection group.**
Difference between the baseline day vs. days post facial paralysis. Significance level =0.05.

**Table 3-3**

| Crush | | | | | | | |
|---|---|---|---|---|---|---|---|
| Chi- square test | | | | | | | |
| | High amplitudes | | | | Low amplitudes | | |
| | p value | percentage | N | | p value | percentage | N |
| .5 hrs | 2.32E-09 | 12.1289228 | 143 | | 9.37E-09 | 87.8710772 | 1036 |
| 6 hrs | 0.00014479 | 30.0469484 | 128 | | 0.00016274 | 69.9530516 | 298 |
| Day 1 | 3.10E-13 | 2.02020202 | 2 | | 2.54E-12 | 97.979798 | 97 |
| Day 2 | 3.10E-13 | 2.02020202 | 2 | | 2.54E-12 | 97.979798 | 97 |
| Day 3 | 1.14E-13 | 1.01010101 | 1 | | 1.01E-12 | 98.989899 | 98 |
| Day 4 | 1.14E-13 | 1.01010101 | 1 | | 1.01E-12 | 98.989899 | 98 |
| Day 5 | 1.13E-13 | 1 | 1 | | 9.96E-13 | 99 | 99 |
| Day 6 | 1.13E-13 | 1 | 1 | | 9.96E-13 | 99 | 99 |
| Day 7 | 1.13E-13 | 1 | 1 | | 9.96E-13 | 99 | 99 |
| Day 8 | 1.14E-13 | 1.01010101 | 1 | | 1.01E-12 | 98.989899 | 98 |
| Day 9 | 1.52E-12 | 3.66972477 | 4 | | 1.11E-11 | 96.3302752 | 105 |
| Day 10 | 1.04E-13 | 0.91743119 | 1 | | 9.22E-13 | 99.0825688 | 108 |
| Day 11 | 1.52E-12 | 3.66972477 | 4 | | 1.11E-11 | 96.3302752 | 105 |
| Day 12 | 8.15E-09 | 13.7614679 | 15 | | 2.94E-08 | 86.2385321 | 94 |
| Day 13 | 9.56E-11 | 8.25688073 | 9 | | 5.08E-10 | 91.7431193 | 100 |
| Day 14 | 3.75E-06 | 22.9357798 | 25 | | 7.06E-06 | 77.0642202 | 84 |
| Day 15 | 1.15E-07 | 17.4311927 | 19 | | 3.19E-07 | 82.5688073 | 90 |
| Day 16 | 4.42E-05 | 27.5229358 | 30 | | 5.98E-05 | 72.4770642 | 79 |
| Day 17 | 2.79E-05 | 26.6055046 | 29 | | 4.03E-05 | 73.3944954 | 80 |
| Day 18 | 2.13E-07 | 18.3486239 | 20 | | 5.57E-07 | 81.6513761 | 89 |
| Day 19 | 4.42E-05 | 27.5229358 | 30 | | 5.98E-05 | 72.4770642 | 79 |
| Day 20 | 0.00010599 | 29.3577982 | 32 | | 0.00012533 | 70.6422018 | 77 |

**Statistical details in the proportion of high and low amplitudes in crush group.** Difference between the baseline day vs. days post facial paralysis. Significance level =0.05.

Table 3-4

| Sham | | | | | | | |
|---|---|---|---|---|---|---|---|
| Chi- square test | | | | | | | |
| | High amplitudes | | | | Low amplitudes | | |
| | p value | percentage | N | | p value | percentage | N |
| .5 hrs | 0.53933387 | 48.0716253 | 698 | | 0.58637691 | 51.9283747 | 754 |
| 6 hrs | 0.02290352 | 59.0909091 | 78 | | 0.38506648 | 62.5 | 330 |
| Day 1 | 0.32712962 | 37.5 | 198 | | 0.04373648 | 40.9090909 | 54 |
| Day 2 | 0.72501583 | 41.6666667 | 55 | | 0.75518889 | 58.3333333 | 77 |
| Day 3 | 0.99271006 | 43.9393939 | 58 | | 0.99353814 | 56.0606061 | 74 |
| Day 4 | 0.08502074 | 32.5757576 | 43 | | 0.12685339 | 67.4242424 | 89 |
| Day 5 | 0.02182996 | 28.7878788 | 38 | | 0.04207221 | 71.2121212 | 94 |
| Day 6 | 0.65437069 | 46.969697 | 62 | | 0.69148364 | 53.030303 | 70 |
| Day 7 | 0.1352144 | 34.0909091 | 45 | | 0.18545059 | 65.9090909 | 87 |
| Day 8 | 3.78E-05 | 16.6666667 | 22 | | 0.00025963 | 83.3333333 | 110 |
| Day 9 | 0.05330847 | 56.8181818 | 75 | | 0.08673064 | 43.1818182 | 57 |
| Day 10 | 0.00210854 | 64.3939394 | 85 | | 0.00642519 | 35.6060606 | 47 |
| Day 11 | 0.17339698 | 53.030303 | 70 | | 0.22753807 | 46.969697 | 62 |
| Day 12 | 0.73876457 | 46.2121212 | 61 | | 0.76753013 | 53.7878788 | 71 |
| Day 13 | 0.57417904 | 47.7272727 | 63 | | 0.61842923 | 52.2727273 | 69 |
| Day 14 | 0.03070837 | 58.3333333 | 77 | | 0.05544505 | 41.6666667 | 55 |
| Day 15 | 0.35610756 | 37.8787879 | 50 | | 0.41336782 | 62.1212121 | 82 |
| Day 16 | 0.30832505 | 50.7575758 | 67 | | 0.36651579 | 49.2424242 | 65 |
| Day 17 | 0.00095684 | 65.9090909 | 87 | | 0.00341451 | 34.0909091 | 45 |
| Day 18 | 0.49896678 | 48.4848485 | 64 | | 0.54896412 | 51.5151515 | 68 |
| Day 19 | 0.06628696 | 31.8181818 | 42 | | 0.10355374 | 68.1818182 | 90 |
| Day 20 | 0.21233863 | 52.2727273 | 69 | | 0.26894746 | 47.7272727 | 63 |

**Statistical details in the proportion of high and low amplitudes in sham group.** Difference between the baseline day vs. days post facial paralysis. Significance level =0.05.

**Table 4-1**

| Transection posterior area | | | | | Transection middle area | | | | Transection anterior area | | |
|---|---|---|---|---|---|---|---|---|---|---|---|
| Analysis: one way ANOVA | | | | | Analysis: one way ANOVA | | | | Analysis: one way ANOVA | | |
| df | F value | p value | | | df | F value | p value | | df | F value | p value |
| 22 | 1.98568404 | 0.02501113 | | | 22 | 3.01086235 | 0.00080586 | | 22 | 5.16887522 | 1.37E-06 |
| Post hoc Tukye | | | | | Post hoc Tukye | | | | Post hoc Tukye | | |
| Comparationlow confidence interval | low confidence inteval | high confidence interval | p value | | low confidence inteval | high confidence interval | p value | | low confidence inteval | high confidence interval | p value |
| .5 hrs | -0.018964 | 1.3953445 | 0.06506211 | | 0.27230543 | 1.4851494 | 0.00026894 | | 0.39573887 | 1.3237196 | 1.29E-06 |
| 6 hrs | -0.0519695 | 1.3623389 | 0.10079956 | | 0.32170647 | 1.5345504 | 9.48E-05 | | 0.1706228 | 1.0986035 | 0.00074793 |
| Day 1 | -0.0756797 | 1.3386288 | 0.13562822 | | 0.28836226 | 1.5012062 | 0.00019199 | | 0.19813761 | 1.1261183 | 0.00035543 |
| Day 2 | 0.06866986 | 1.4829783 | 0.01811275 | | 0.31953555 | 1.5323794 | 9.92E-05 | | 0.45431277 | 1.3822935 | 1.55E-07 |
| Day 3 | 0.02064103 | 1.4349494 | 0.03720494 | | 0.29330754 | 1.5061514 | 0.00017298 | | 0.46798274 | 1.3959634 | 7.01E-08 |
| Day 4 | -0.0625867 | 1.3517218 | 0.1153489 | | 0.13537121 | 1.3482151 | 0.00431938 | | 0.47751591 | 1.4054966 | 2.71E-08 |
| Day 5 | -0.152395 | 1.2619134 | 0.31449121 | | 0.2667529 | 1.4795969 | 0.00030207 | | 0.33906248 | 1.2670432 | 6.89E-06 |
| Day 6 | -0.2152659 | 1.1990426 | 0.53312689 | | 0.1583572 | 1.371201 | 0.00275307 | | 0.45332471 | 1.3813055 | 1.63E-07 |
| Day 7 | 0.06507176 | 1.4793801 | 0.01914364 | | 0.13151646 | 1.3443604 | 0.00465455 | | 0.38073978 | 1.3087205 | 2.04E-06 |
| Day 8 | 0.10954159 | 1.52385 | 0.0095207 | | 0.27245194 | 1.4852958 | 0.00026815 | | 0.36002603 | 1.2880068 | 3.76E-06 |
| Day 9 | 0.03835154 | 1.45266 | 0.02867355 | | 0.34089684 | 1.5537407 | 6.30E-05 | | 0.35821596 | 1.2861967 | 3.96E-06 |
| Day 10 | 0.0938822 | 1.5081906 | 0.01221775 | | 0.18920183 | 1.4020457 | 0.00148738 | | 0.43966916 | 1.3676499 | 2.94E-07 |

| | | | | | | | | | | |
|---|---|---|---|---|---|---|---|---|---|---|
| Day 11 | 0.00172216 | 1.4160306 | 0.04879763 | | 0.13790977 | 1.3507537 | 0.00411144 | | 0.43063542 | 1.3586161 | 4.10E-07 |
| Day 12 | -0.118279 | 1.2960293 | 0.22166331 | | 0.24242646 | 1.4552703 | 0.00050109 | | 0.274905 | 1.2028857 | 4.25E-05 |
| Day 13 | -0.3050938 | 1.1092145 | 0.84492576 | | 0.15158343 | 1.3644273 | 0.00314647 | | 0.29383722 | 1.221818 | 2.49E-05 |
| Day 14 | 0.20779371 | 1.6221021 | 0.00185547 | | 0.04883152 | 1.2616754 | 0.02167619 | | 0.50240517 | 1.4303858 | -4.32E-08 |
| Day 15 | 0.01864469 | 1.4329531 | 0.03829856 | | 0.01077449 | 1.2236184 | 0.04181074 | | 0.33957621 | 1.2675569 | 6.79E-06 |
| Day 16 | -0.1291753 | 1.2851331 | 0.2489731 | | 0.19001663 | 1.4028605 | 0.00146319 | | 0.39186338 | 1.3198441 | 1.46E-06 |
| Day 17 | 0.10062999 | 1.5149384 | 0.01097753 | | 0.11361337 | 1.3264573 | 0.00656569 | | 0.40859351 | 1.3365742 | 8.66E-07 |
| Day 18 | -0.0468044 | 1.3675041 | 0.09429703 | | 0.21125758 | 1.4241015 | 0.00095097 | | 0.30926922 | 1.23725 | 1.61E-05 |
| Day 19 | 0.00905287 | 1.4233613 | 0.04396841 | | -0.015891 | 1.1969529 | 0.0646777 | | 0.34196219 | 1.2699429 | 6.33E-06 |
| Day 20 | 0.03635639 | 1.4506648 | 0.02953629 | | 0.10167348 | 1.3145174 | 0.00823355 | | 0.4378148 | 1.3657955 | 3.15E-07 |

**Statistical details in the differences between frames in transection group.** Difference between the first frame with the others in the video, comparation between baseline vs days post facial paralysis. Significance level =0.05.

**Table 4-2**

| | | | | | | | | | | | |
|---|---|---|---|---|---|---|---|---|---|---|---|
| Crush posterior area | | | | | Crush middle area | | | | Crush anterior area | | |
| Analysis: one way ANOVA | | | | | Analysis: one way ANOVA | | | | Analysis: one way ANOVA | | |
| df | F value | p value | | | df | F value | p value | | df | F value | p value |
| 22 | 2.12548065 | 0.01560637 | | | 22 | 3.76091599 | 7.52E-05 | | 22 | 4.72901535 | 4.50E-06 |
| Post hoc Tukye | | | | | Post hoc Tukye | | | | Post hoc Tukye | | |
| Comparationlow confidence interval | low confidence inteval | high confidence interval | p value | | low confidence inteval | high confidence interval | p value | | low confidence inteval | high confidence interval | p value |
| .5 hrs | -0.5589215 | 1.0581894 | 0.99985915 | | 0.36278164 | 1.5345566 | 3.00E-05 | | 0.08144373 | 1.3261409 | 0.01247929 |
| 6 hrs | -0.1639256 | 1.4531853 | 0.28785354 | | 0.32556146 | 1.4973364 | 6.85E-05 | | 0.20385897 | 1.4485561 | 0.00122909 |
| Day 1 | -0.3247579 | 1.2923529 | 0.78269285 | | 0.32208389 | 1.4938588 | 7.40E-05 | | 0.32631481 | 1.5710119 | 0.00010264 |
| Day 2 | -0.0451583 | 1.5719526 | 0.08570348 | | 0.26873499 | 1.44051 | 0.00023797 | | 0.23976284 | 1.4844599 | 0.00060093 |
| Day 3 | -0.2585768 | 1.3585341 | 0.57443261 | | 0.3022337 | 1.4740087 | 0.00011449 | | 0.24808675 | 1.4927838 | 0.00050832 |
| Day 4 | -0.5283838 | 1.088727 | 0.99923044 | | 0.28739589 | 1.4591708 | 0.0001584 | | 0.35177535 | 1.5964725 | 6.05E-05 |
| Day 5 | -0.2451534 | 1.3719575 | 0.52988642 | | 0.34810841 | 1.5198834 | 4.15E-05 | | 0.22293895 | 1.4676361 | 0.00084166 |
| Day 6 | -0.1841391 | 1.4329717 | 0.34135166 | | 0.29827052 | 1.4700456 | 0.00012487 | | 0.29208148 | 1.5367786 | 0.00020783 |
| Day 7 | -0.0282502 | 1.5888608 | 0.0703482 | | 0.27919728 | 1.4509723 | 0.00018951 | | 0.23399103 | 1.4786881 | 0.00067479 |
| Day 8 | -0.3149961 | 1.3021147 | 0.75473726 | | 0.32449281 | 1.4962678 | 7.01E-05 | | 0.25244755 | 1.4971447 | 0.00046547 |
| Day 9 | -0.0698816 | 1.5472293 | 0.11323776 | | 0.22321588 | 1.3949909 | 0.00063397 | | 0.26362765 | 1.5083247 | 0.00037116 |
| Day 10 | -0.6981545 | 0.91895634 | 0.99999875 | | 0.1967591 | 1.3685341 | 0.00111015 | | -0.2323326 | 1.0123645 | 0.71505177 |

| | | | | | | | | | | | |
|---|---|---|---|---|---|---|---|---|---|---|---|
| Day 11 | -0.1100537 | 1.5070572 | 0.17316373 | | 0.17030233 | 1.3420773 | 0.00192822 | | -0.2172025 | 1.0274945 | 0.65244621 |
| Day 12 | 0.08740407 | 1.704515 | 0.0160597 | | 0.2402696 | 1.4120445 | 0.00044012 | | -0.0456387 | 1.1990583 | 0.1006551 |
| Day 13 | -0.2775111 | 1.3395998 | 0.63714176 | | 0.17071962 | 1.3424946 | 0.0019117 | | -0.1807303 | 1.0639668 | 0.49590263 |
| Day 14 | -0.3738774 | 1.2432334 | 0.89777726 | | 0.13132381 | 1.3030988 | 0.00427305 | | -0.2501896 | 0.99450749 | 0.78349721 |
| Day 15 | -0.2611921 | 1.3559188 | 0.58313072 | | 0.06326246 | 1.2350374 | 0.01608122 | | -0.1844164 | 1.0602807 | 0.51157218 |
| Day 16 | -0.2477484 | 1.3693624 | 0.5384711 | | 0.02676553 | 1.1985404 | 0.03136139 | | 0.0128715 | 1.2575686 | 0.04058885 |
| Day 17 | -0.4555275 | 1.1615833 | 0.98577821 | | -0.0097314 | 1.1620436 | 0.05891329 | | -0.1489977 | 1.0956994 | 0.36813438 |
| Day 18 | -0.3889422 | 1.2281687 | 0.92333037 | | -0.0887544 | 1.0830207 | 0.19599441 | | -0.1461186 | 1.0985785 | 0.35739094 |
| Day 19 | -0.1789375 | 1.4381733 | 0.32706818 | | -0.0080537 | 1.1637213 | 0.05728279 | | -0.0361619 | 1.2085352 | 0.087519 |
| Day 20 | -0.1164183 | 1.5006926 | 0.1845856 | | 0.0623169 | 1.2340919 | 0.01636869 | | 0.07497919 | 1.3196763 | 0.01400852 |

**Statistical details in the differences between frames in crush group.** Difference between the first frame with the others in the video, comparation between baseline vs days post facial paralysis. Significance level =0.05.

**Table 4-3**

| Sham posterior area | | | | | Sham middle area | | | | Sham anterior area | | |
|---|---|---|---|---|---|---|---|---|---|---|---|
| Analysis: one way ANOVA | | | | | Analysis: one way ANOVA | | | | Analysis: one way ANOVA | | |
| df | F value | p value | | | df | F value | p value | | df | F value | p value |
| 22 | 0.44879752 | 0.97764826 | | | 22 | 1.54423678 | 0.10641832 | | 22 | 2.19553185 | 0.0123127 |
| Post hoc Tukye | | | | | Post hoc Tukye | | | | Post hoc Tukye | | |
| Comparationlow confidence interval | low confidence inteval | high confidence interval | p value | | low confidence inteval | high confidence interval | p value | | low confidence inteval | high confidence interval | p value |
| .5 hrs | -1.0951746 | 0.6982885 | 0.99999827 | | -0.7443456 | 0.81382954 | 0.99999875 | | -0.787832 | 0.78234756 | 0.99999875 |
| 6 hrs | -0.8135519 | 0.97991121 | 0.99999875 | | -0.9208729 | 0.63730216 | 0.99999875 | | -0.7956059 | 0.77457368 | 0.99999875 |
| Day 1 | -0.9362071 | 0.857256 | 0.99999875 | | -0.9883462 | 0.56982887 | 0.99998391 | | -1.0234889 | 0.5466907 | 0.9998908 |
| Day 2 | -0.8205403 | 0.9729228 | 0.99999875 | | -0.8738245 | 0.68435061 | 0.99999875 | | -0.9089946 | 0.66118503 | 0.99999875 |
| Day 3 | -1.0024329 | 0.79103017 | 0.99999875 | | -0.7636162 | 0.79455888 | 0.99999875 | | -0.9159247 | 0.65425491 | 0.99999875 |
| Day 4 | -1.08132 | 0.712143 | 0.99999863 | | -0.9329022 | 0.62527293 | 0.99999869 | | -0.6968494 | 0.87333024 | 0.99999875 |
| Day 5 | -0.7557613 | 1.0377018 | 0.99999875 | | -1.0035428 | 0.55463237 | 0.99995184 | | -0.8268335 | 0.7433461 | 0.99999875 |
| Day 6 | -0.9121696 | 0.88129348 | 0.99999875 | | -0.9547927 | 0.60338235 | 0.99999809 | | -0.959147 | 0.61103261 | 0.99999821 |
| Day 7 | -0.8213451 | 0.97211802 | 0.99999875 | | -1.2257459 | 0.33242923 | 0.8357116 | | -0.6340872 | 0.93609238 | 0.99999869 |
| Day 8 | -0.9110919 | 0.88237125 | 0.99999875 | | -0.817365 | 0.7408101 | 0.99999875 | | -0.9686954 | 0.60148418 | 0.99999738 |
| Day 9 | -0.9415035 | 0.85195965 | 0.99999875 | | -0.9288681 | 0.62930697 | 0.99999869 | | -0.8084906 | 0.76168895 | 0.99999875 |
| Day 10 | -1.085415 | 0.70804811 | 0.99999857 | | -1.1714947 | 0.38668036 | 0.94082814 | | -0.9846116 | 0.58556801 | 0.99999297 |

| | | | | | | | | | | | |
|---|---|---|---|---|---|---|---|---|---|---|---|
| Day 11 | -0.9212912 | 0.87217194 | 0.99999875 | | -1.1300827 | 0.42809236 | 0.98004013 | | -0.9870934 | 0.58308619 | 0.9999916 |
| Day 12 | -0.92792 | 0.86554313 | 0.99999875 | | -1.0338666 | 0.52430844 | 0.9996658 | | -1.1847303 | 0.38544923 | 0.93504006 |
| Day 13 | -0.9780067 | 0.81545639 | 0.99999875 | | -1.0628071 | 0.495368 | 0.99847245 | | -0.8726939 | 0.69748569 | 0.99999875 |
| Day 14 | -0.7969792 | 0.99648392 | 0.99999875 | | -1.1832244 | 0.37495071 | 0.92362344 | | -0.9160974 | 0.65408218 | 0.99999875 |
| Day 15 | -0.9790111 | 0.81445205 | 0.99999875 | | -0.753053 | 0.80512214 | 0.99999875 | | -0.9150201 | 0.65515947 | 0.99999875 |
| Day 16 | -0.7794061 | 1.0140569 | 0.99999875 | | -1.115078 | 0.44309711 | 0.98760289 | | -1.2583408 | 0.31183881 | 0.77255231 |
| Day 17 | -1.1229515 | 0.67051154 | 0.99999368 | | -1.2866516 | 0.27152342 | 0.65115482 | | -1.2162427 | 0.35393691 | 0.87951624 |
| Day 18 | -0.8742496 | 0.91921347 | 0.99999875 | | -1.3377128 | 0.22046238 | 0.47643399 | | -1.3639631 | 0.20621645 | 0.42449856 |
| Day 19 | -0.7459262 | 1.0475368 | 0.99999875 | | -0.7697285 | 0.78844655 | 0.99999875 | | -0.6055963 | 0.96458328 | 0.99999785 |
| Day 20 | -0.9814837 | 0.81197941 | 0.99999875 | | -1.0629236 | 0.49525154 | 0.99846399 | | -1.4006104 | 0.16956913 | 0.31552583 |

**Statistical details in the differences between frames in sham group.** Difference between the first frame with the others in the video, comparation between baseline vs days post facial paralysis. Significance level =0.05.

**Table 6-1**

| T-test | | | |
|---|---|---|---|
| facial palsy model | df | sd value | p value |
| sucrose vs pleasure | 899 | 0.18200533 | 5.60E-86 |
| quinine vs disgust | 899 | 0.2324 | 1.20E-27 |
| water vs neutral | 899 | 0.2074 | 5.21E-50 |

**Statistical details in facial expression before and after oral stimulation with solutions.** Comparation between 10 seconds pre and post stimulation with sucrose, quinine or water using the pleasure, disgust and neutral prototype. Significance level =0.05.

**Table 6-2**

| T-test | | | |
|---|---|---|---|
| Comparation | sd | df value | p value |
| Transection day 1 | 0.208 | 899 | 1E-31 |
| Transection day 20 | 0.1878 | 899 | 1.00E-36 |
| Crush day 1 | 0.2151 | 899 | 3.05E-13 |
| Crush day 20 | 0.1721 | 899 | 1.00E-36 |

**Statistical details in facial expression between baseline vs facial paralysis.** Similarity of pleasure prototype for 10 second after oral stimulation with sucrose between baseline, day 1 and day 20 post facial paralysis in transection and crush group. Significance level =0.05.

**Table 7-1**

| Wiloxon signed rank-tets | | | | |
|---|---|---|---|---|
| | z value | p value | N of mice | N of neurons |
| Transection baseline | -3.7642 | 1.67E-04 | 2 | 21, mouse1= 12, mouse2= 9 |
| transection day 1 | 1.5722 | 0.11591433 | 2 | 20, mouse1= 11, mouse2= 9 |
| transection day 20 | -1.2612 | 0.20723854 | 2 | 17, mouse1= 9, mouse2= 8 |
| Crush baseline | 8.4251 | 3.60E-17 | 2 | 24, mouse1= 12, mouse2= 12 |
| Crush day 1 | -0.6008 | 0.54799886 | 2 | 22, mouse1= 12, mouse2= 10 |
| Crush day 20 | -11.6501 | 2.29E-31 | 2 | 15, mouse1= 9, mouse2= 6 |

**Statistical details in population neuronal activity in ALM pre and post facial expression.** Differences between z-score pre and post oral stimulation with sucrose in population neuronal activity of two mice. Significance level =0.05.

**Table 7-2**

| Wiloxon signed rank-tets | | | | | | | | |
|---|---|---|---|---|---|---|---|---|
| Mouse walking | | | | Stepper motor | | | | |
| | z value | p value | | | z value | p value | N of mice | N of neurons |
| Transection basal | 1.5335 | 1.25E-01 | | Transection basal | -2.2644 | 5.36E-02 | 2 | 21, mouse1= 12, mouse2= 9 |
| transection day 1 | 1.4544 | 0.14583228 | | transection day 1 | 8.4497 | 0.092 | 2 | 20, mouse1= 11, mouse2= 9 |
| transection day 20 | -2.1696 | 0.06003845 | | transection day 20 | -4.9134 | 8.90E-02 | 2 | 17, mouse1= 9, mouse2= 8 |
| Crush basal | 0.4264 | 0.66985117 | | Crush basal | -0.332 | 7.40E-01 | 2 | 24, mouse1= 12, mouse2= 12 |
| Crush day 1 | 1.4544 | 0.25 | | Crush day 1 | -0.5536 | 5.80E-01 | 2 | 22, mouse1= 12, mouse2= 10 |
| Crush day 20 | -7.1483 | 8.74E-02 | | Crush day 20 | 2.4611 | 5.39E-02 | 2 | 15, mouse1= 9, mouse2= 6 |

**Statistical details in population neuronal activity in ALM pre and post control events.** Differences between z-score pre and post walking and activation of stepper motor in population neuronal activity of two mice. Significance level =0.05.

**Extended Data 1**

Extended data includes a copy of the GitHub repository. Contains a copy of code with two sections. Section 1: code to develop FaPDA with any video recording information of facial paralysis. Section 2: code to use FaPDA in video recordings of mice with facial paralysis.

```
%% Threshold calculation

clc, clear all, close all %% Removes system variables and allows the program to start

cd 'C:\Users\warri\Dropbox\' %% Identify the address where the videos to be used are located

video_baseline=VideoReader('R1transA.mp4');%% reading the video to be used of baseline

video_transection=VideoReader('R1transA.mp4');%% reading the video to be used of transection day 1

  N= video_baseline.NumberOfFrames;%% number of frames in baseline video

  n= video_transection.NumberOfFrames;%% number of frames in transction video

load('coordinates.mat')%% loading of anterior and middle facial region coordinates

HOGs=[]; %Create a new variable

HOGs2=[];%Create a new variable

for loop=1:N %% loop for extract HOGs for video

  image=read(video_baseline,loop);%% Extract frame to frame from the video

  imageanterior=imcrop(image, coordenadas(1,:));%% image cut of the anterior area of the face

  imagemiddle=imcrop(image, coordenadas(1,:));%% image cut of the middle area of the face
```

```
  imageanteriorgray= rgb2gray(croppedImg);%% transform image from RGB to grayscale

  imagemiddlegray= rgb2gray(croppedImg);%% transform image from RGB to grayscale

  [featureVector,hogVisualization] = extractHOGFeatures(imageanteriorgray, 'CellSize', [32 32],'BlockSize',[1 1] ,'NumBins',8); %% Extract HOGs fetures from cut image of anterior area

  [featureVector2,hogVisualization2] = extractHOGFeatures(imagemiddlegray, 'CellSize', [32 32],'BlockSize',[1 1] ,'NumBins',8);%% Extract HOGs fetures from cut image of middle area

   HOGs=[HOGs;featureVector];% Save HOGs features in variable HOGs

   HOGs2=[HOGs2;featureVector2];%Save HOGs features in variable HOGs2

end

for loop=1:length(HOGs)% loop to obtain differences between frames

   difference(loop)=HOGs(1)-HOGs(loop)%Creat a variable with differences between frame 1 and all the frames in anterior area of baseline video

   difference2(loop)=HOGs2(1)-HOGs2(loop)%Creat a variable with differences between frame 1 and all the frames in middle area of baseline video

end

HOGs=[]; %Create a new variable

HOGs2=[];%Create a new variable

for loop=1:N %% loop for extract HOGs for video

  image=read(video_transection,loop);%% Extract frame to frame from the video
```

```
   imageanterior=imcrop(image, coordenadas(1,:));%% image cut of the anterior area of the face

   imagemiddle=imcrop(image, coordenadas(1,:));%% image cut of the middle area of the face

   imageanteriorgray= rgb2gray(croppedImg);%% transform image from RGB to grayscale

   imagemiddlegray= rgb2gray(croppedImg);%% transform image from RGB to grayscale

   [featureVector,hogVisualization] = extractHOGFeatures(imageanteriorgray, 'CellSize', [32 32],'BlockSize',[1 1] ,'NumBins',8); %% Extract HOGs fetures from cut image of anterior area

   [featureVector2,hogVisualization2] = extractHOGFeatures(imagemiddlegray, 'CellSize', [32 32],'BlockSize',[1 1] ,'NumBins',8);%% Extract HOGs fetures from cut image of middle area

    HOGs=[HOGs;featureVector];% Save HOGs features in variable HOGs

    HOGs2=[HOGs2;featureVector2];%Save HOGs features in variable HOGs2

end

for loop=1:length(HOGs)% loop to obtain differences between frames

    difference3(loop)=HOGs(1)-HOGs(loop)%Creat a variable with differences between frame 1 and all the frames in anterior area of transection video

    difference4(loop)=HOGs2(1)-HOGs2(loop)%Creat a variable with differences between frame 1 and all the frames in middle area of transection video

end
```

thersholdanterior=(mean(difference)+mean(difference3))./2% Creat a thershold of anterior area

thersholdmiddle=(mean(difference2)+mean(difference4))./2% Creat a thershold of middle area

%% Use FaPDA: Facial paralysis detection algorithm applied in mice

clc, clear all, close all %% Removes system variables and allows the program to start

cd 'C:\Users\warri\Dropbox\' %% Identify the address where the videos to be used are located

for loop=1:23 %% Loop with all the evaluated videos

  video=VideoReader(['R1trans' num2str(loop) '.mp4']);%% reading the video to be used

HOGs=[]; %Create a new variable

HOGs2=[];%Create a new variable

for loop2=1:N %% loop for extract HOGs for video

  image=read(video,loop2);%% Extract frame to frame from the video

  imageanterior=imcrop(image, coordenadas(1,:));%% image cut of the anterior area of the face

  imagemiddle=imcrop(image, coordenadas(1,:));%% image cut of the middle area of the face

  imageanteriorgray= rgb2gray(croppedImg);%% transform image from RGB to grayscale

  imagemiddlegray= rgb2gray(croppedImg);%% transform image from RGB to grayscale

```
   [featureVector,hogVisualization] = extractHOGFeatures(imageanteriorgray, 'CellSize', [32 32],'BlockSize',[1 1] ,'NumBins',8); %% Extract HOGs fetures from cut image of anterior area

   [featureVector2,hogVisualization2] = extractHOGFeatures(imagemiddlegray, 'CellSize', [32 32],'BlockSize',[1 1] ,'NumBins',8);%% Extract HOGs fetures from cut image of middle area

   HOGs=[HOGs;featureVector];% Save HOGs features in variable HOGs

   HOGs2=[HOGs2;featureVector2];%Save HOGs features in variable HOGs2

end

for loop3=1:length(HOGs)% loop to obtain differences between frames

   difference1=HOGs(1)-HOGs(loop3)%Creat a variable with differences between frame 1 and all the frames in anterior area of video

   difference2=HOGs2(1)-HOGs2(loop3)%Creat a variable with differences between frame 1 and all the frames in middle area of video

if difference1<thersholdanterior %conditional to detect frames with values under the thershold

   identityanterior(loop3)=1% frame without movement

else

   identityanterior(loop3)=0% frame with movement

end

if difference1<thersholdanterior %conditional to detect frames with values under the thershold

   identitymiddle(loop3)=1% frame without movement

else

   identitymiddle(loop3)=0% frame with movement

end

end
```

```
if length(find(identityanterior==1))>(95*N)/100 &
length(find(identitymiddle==1))>(95*N)/100 %Detection conditional

    detection(loop)=1%If 95% of the frames are without movement, the system ends up
in paralysis.

else

    detection(loop)=0%If 95% of the frames are without movement, the system ends up
in without paralysis

end

end
```
